# Large-Scale Labeling and Assessment of Sex Bias in Publicly Available Expression Data

**DOI:** 10.1101/2020.10.26.356287

**Authors:** Emily Flynn, Annie Chang, Russ B. Altman

## Abstract

Women are at more than 1.5-fold higher risk for clinically relevant adverse drug events. While this higher prevalence is partially due to gender-related effects, biological sex differences likely also impact drug response. Publicly available gene expression databases provide a unique opportunity for examining drug response at a cellular level. However, missingness and heterogeneity of metadata prevent large-scale identification of drug exposure studies and limit assessments of sex bias. To address this, we trained organism-specific models to infer sample sex from gene expression data, and used entity normalization to map metadata cell line and drug mentions to existing ontologies. Using this method, we infer sex labels for 450,371 human and 245,107 mouse microarray and RNA-seq samples from refine.bio. Overall, we find slight female bias (52.1%) in human samples and (62.5%) male bias in mouse samples; this corresponds to a majority of single sex studies, split between female-only and male-only (33.3% vs 18.4% in human and 31.0% vs 30.4% in mouse respectively). In drug studies, we find limited evidence for sex-sampling bias overall; however, specific categories of drugs, including human cancer and mouse nervous system drugs, are enriched in female-only and male-only studies respectively. Our expression-based sex labels allow us to further examine the complexity of cell line sex and assess the frequency of metadata sex label misannotations (2-5%). We make our inferred and normalized labels, along with flags for misannotated samples, publicly available to catalyze the routine use of sex as a study variable in future analyses.

## INTRODUCTION

Sex differences have been reported across multiple traits and diseases and in response to drugs. In the case of drug response, women experience more than 1.5-fold as many adverse drug events (Zopf et al. 2008). This is in part due to historical exclusion of women from clinical research. In 1993, the policies excluding women were revoked and the National Institutes of Health (NIH) Revitalization Act was passed to increase inclusion of women and minorities in clinical research. This has improved inclusion of women, but clinical studies continue to show sex bias against female participants (Kim, Tingen, and Woodruff 2010; Prakash et al. 2018; Feldman et al. 2019). Additionally, preclinical studies are critical to the drug development process (Tannenbaum, Day, and Matera Alliance 2017); however, there is limited reporting of sex in both rodent (Beery and Zucker 2011; Klein et al. 2015) and cell line research (Shah, McCormack, and Bradbury 2014). In 2016, the NIH passed a mandate that requires researchers to consider sex as a variable in preclinical analysis (Clayton and Collins 2014), and this has led to increases in sex reporting, but sex bias in these studies still remains (Woitowich, Beery, and Woodruff 2020).

Gene expression data is often used as part of the drug development pipeline in order to better understand cellular and molecular-level effects of drugs and assess their mechanisms of action and side effects (Chengalvala et al. 2007). While we do not expect all drugs to show cellular-level sex differences in drug response, pervasive use of single-sex studies may lead to the development of drugs that do not work well for both men and women.

Multiple studies have sought to assess sex reporting and bias in specific areas, including in skin (Kong et al. 2016), neuroscience (Mamlouk et al. 2020; Beery and Zucker 2011), and pain research (Mogil and Chanda 2005), and across biomedical fields (Beery and Zucker 2011; Woitowich, Beery, and Woodruff 2020); these assessments largely focus on scientific literature. Public repositories of biological data provide another avenue for assessing sex bias. Repositories such as Gene Expression Omnibus (GEO) (Edgar, Domrachev, and Lash 2002), Sequence Read Archive (SRA)(Leinonen et al. 2010), and ArrayExpress (Brazma et al. 2003) contain gene expression data and corresponding metadata describing the experiments and samples, allowing for re-analysis and re-use of these data. Gene expression metadata is organized according to the Minimum Information about a Microarray Experiment (MIAME)(Brazma et al. 2001) and Minimum Information about a high-throughput Nucleotide Sequencing Experiment (MINISEQE) guidelines; despite this, metadata is heterogeneous across experiments, making large-scale analysis difficult. In the case of examining sex labels, not only are these labels reported inconsistently, the majority of samples are missing this information, limiting assessment of sex bias. Previous studies also suggest there may also be widespread misannotation (Toker, Feng, and Pavlidis 2016; Lohr et al. 2015). Metadata normalization, such as that performed by MetaSRA (Bernstein, Doan, and Dewey 2017), seeks to address the problem of inconsistent reporting of sex, cell type, age, and multiple other labels by mapping these to existing ontologies; however, it is unable to address the issue of missingness due to lack of metadata.

Assessment of sex bias in gene expression data does not require metadata: sex can be imputed from the expression levels of X and Y chromosome genes. Recount2 (Collado-Torres et al. 2017) used expression data to impute sex labels across 70,000 publicly available human RNA-seq samples using a linear model and found slight female bias overall; however their work does not extend to microarray platforms (Ellis et al. 2018). While other methods for imputing sex labels from microarray expression data have good performance, they are either clustering based and therefore limited to mixed sex studies (because they assume two clusters) (Buckberry et al. 2014; Toker, Feng, and Pavlidis 2016) or only work for specific platforms (e.g., (Giles et al. 2017) GPL570).

In addition to metadata challenges, expression data can be difficult to study at scale because of platform heterogeneity and differences in data pre-processing and normalization (Ramasamy et al. 2008). Previous resources, such as ARCHS4 (Lachmann et al. 2018) and recount2 (Collado-Torres et al. 2017) have worked to address these limitations by releasing human and mouse RNA-seq data that has been processed with standardized pipelines.

Refine.bio is a new transcriptomic resource that addresses many of these previous limitations (http://www.refine.bio, Greene et al.). It contains all publicly available microarray and RNA-seq data from twenty-two organisms, pre-processed, de-duplicated, and normalized to allow for examination across platforms. Additionally, portions of the metadata are “harmonized”, meaning that groups of semantically similar labels (eg., “tissue” and “organ”) have been manually aggregated into categories.

We sought to assess sex bias at the sample and study level across all publicly available expression data, including specifically drug-related datasets. For this analysis, we focus on human and mouse expression studies, using refine.bio as a resource, and expand on previous analysis by using a penalized logistic regression model to predict sample sex. In the process we also consider cell line sex and use our imputed sex labels to estimate baseline misannotation rates.

## METHODS

The code used for this analysis is publicly available at https://github.com/erflynn/sl_label; all analyses were performed in R (3.6.1) or python (3.7.1);

### 1 Dataset construction

#### 1-1 Expression data pre-processing

We downloaded the normalized expression compendia and RNA-seq libraries for human (*Homo sapiens*) and mouse (*Mus musculus*) from refine.bio (3/15/2020). The compendia contain both microarray and RNA-seq data, quantile normalized and with missing values imputed using SVD-impute (Perry 2009) (430,119 human, 228,708 mouse samples). We extracted the microarray data (330,508 human, 123,279 mouse samples) and converted the data to gctx format to aid analysis (Enache et al. 2019); no other transformations were applied to them. All RNA-seq libraries for human and mouse were downloaded from refine.bio (122,864 human, 125,652 mouse). RNA-seq samples with fewer than 100,000 counts were removed, resulting in 119,863 human and 121,828 mouse samples. Transcripts per million (TPM) counts were extracted from the salmon quant.sf files output (Patro et al. 2017) To convert RNA-seq count data to a normal distribution for logistic regression, the data were transformed with the Box-Cox transformation using the R package BestNormalize (Peterson and Cavanaugh 2019).

#### 1-2 Extraction of sample and study metadata labels

GEO microarray metadata were extracted from GEOMetadb (Zhu et al. 2008), including study information (title, description, date), sample-study membership, and sample titles and attributes included in the *characteristics_ch1* field. Following a similar process to refine-bio metadata harmonization, RNA-seq metadata was extracted from European Nucleotide Archive (Leinonen et al. 2011) XML files, including study information, study-run membership, run-to-sample mappings, and sample attributes. All runs were assigned the attributes of their corresponding samples (each sample can map to multiple runs but no run maps to multiple samples; for consistency with other data sources, we refer to SRA runs as “samples” from here forward). ArrayExpress microarray metadata was also extracted from ENA json files.

Sample metadata was present for 443,611 of 448,827 (98.8%) GEO samples, all 4960 ArrayExpress samples, and 237,267 of 241,691 RNA-seq runs (98.2%, corresponding to 196,852 unique samples).

### 2 Sex labeling

#### 2-1 Metadata sex label extraction

Sample-level sex labels were extracted from metadata sample attributes by filtering for keys that contained the words “sex” or “gender”. Additional attributes were also extracted if values contained exact matches to the words “male” or “female” All unique values were then mapped to one of “male”, “female”, “mixed” sex (e.g. pooled sample from both males and females), or “unlabeled”. The human labels for the refine-bio RNA-seq samples almost exactly match those of MetaSRA (Bernstein, Doan, and Dewey 2017) (30,063 of 30,073 samples).

We grouped studies into the following categories based on the provided sample sex labels:

1. *Unlabeled*: studies with either less than half of their samples labeled (for studies with up to sixty samples) or less than thirty samples labeled (for studies with more than sixty samples)
2. *Male-only*: all male labels
3. *Female-only*: all female labels
4. *Mostly-male*: >80% of labeled samples are male
5. *Mostly-female*: >80% of labeled samples are female
6. *Mixed sex*: ≤80% of labeled samples belong to either sex

To distinguish between studies with similar and highly imbalanced male/female proportions, we created separate categories for mixed sex (≤80%) vs mostly-male and mostly-female studies (>80%). See Supplementary Tables S1A and S1B for the sample and study sex breakdowns respectively.

#### 2-2 Inferring sample sex from expression data

##### 2-2-1 Training and testing data

There is often substantial overlap of samples across studies; with groups of samples belonging to multiple studies. To reduce this overlap, we removed studies that share one or more samples with greater than 5 other studies. For the remainder of studies, we aggregated studies into study groups such that all samples that share any study are in the same study group.

For each organism and data type, we stratified by study group, sampled a maximum of five samples per study (to limit overfitting to a particular study), and then randomly selected training (n=2,300-3,200 samples, 540-1,300 studies) and testing data (n=630-790 samples, 120-360 studies) such that was approximately balanced between males and females (48.4-50.7%) (see Supplementary Table S2A for size of datasets by organism and data type). The goals in constructing these data sets were to ensure there was no leakage between training and test sets, limit overfitting, and retain sufficient samples and studies to perform stratified cross-validation.

##### 2-2-2 Model training

We trained logistic regression models with an elastic net penalty (using the R package glmnet (Friedman, Hastie, and Tibshirani 2010)) using all X and Y chromosome genes as input. We used nested six-fold cross-validation to select the hyperparameters alpha (elastic net penalty) and lambda (shrinkage parameter). Briefly, for each of the six cross-validation folds (study-stratified), each value of alpha (0.1 to 1 in increments of 0.1), lambda was selected from a grid of values by performing cross-validation on five of the six folds and selecting the lambda within one standard error of the minimum mean cross-validated error (“lambda-1se”). This model was assessed on the sixth “validation” fold. We then computed the median classification error and median lambda for each alpha across all six validation folds, and selected the value of alpha (and its corresponding lambda) with the lowest median error.

We performed nested cross-validation in this way because of the substantial between-study heterogeneity and within-study correlations in expression data. Selection of hyperparameters without the additional cross-validation loop led to increased classification error. The procedure described is equivalent to the percentile lasso (Roberts and Nowak 2014) using the 50th percentile, and extended to select both alpha and lambda.

##### 2-2-3 Accuracy assessment and cutoff selection

We assessed the accuracy of our model using both the held out test set (described above) and an extended test set consisting of all samples with metadata sex labels from studies not in either the training set. We consider a sample to be correctly labeled when metadata sex matches the expression-based sex; however, it is important to note that we are actually measuring concordance, as metadata sex labels may contain errors. We also examined two subsets of the extended test set: samples from large (1) *mixed sex* studies (at least 10 samples per study) or (2) *exclusively single sex* studies (at least 8 samples per study, with all samples present and annotated as from one sex) and assessed the accuracy on these subsets of the data (see Supplementary Table S2A for a full list of assessment datasets and their size, and Supplementary Table S2B for the corresponding accuracies).

We created the exclusively single sex datasets for two reasons: (1) we expect that if metadata indicates all samples in a study are of one sex it is less likely there are misannotation errors, and (2) to make sure that within study variability did not skew or ability to label these data. In microarray data in particular because these data are signal intensities rather than counts, improper normalization of single sex studies can lead to a wider distribution of sex relevant gene expression values.

For assigning sex labels to samples, we used a cutoff threshold of 0.7 on the model predicted probabilities in order to approximate 95% accuracy (see Supplementary Figure 2 and Supplementary Table S2B). This leads to labeling of 91-93% of the microarray and 70-73% of the RNA-seq extended test sets.

##### 2-2-4 Assessment of performance across platforms

Platform heterogeneity presents a huge challenge for examining microarray data, and as a result, previous sex labeling methods have been limited to specific platforms. With our method, we aimed to have high performance across a range of platforms. Our models show high performance in the majority of platforms; however, it performs particularly poorly in a small number of platforms (all less than 50% accuracy; no other platforms have between 50-70% accuracy) but these cover <3% of samples (Supplementary Figure 3 and Supplementary Table 3A-B). As a result, these six platforms were excluded from subsequent analysis.

### 3. Cell line sex label assessment

#### 3-1 Cell line labeling

##### 3-1-1 Cell line normalization

Cell line names, synonyms, sex labels, and amelogenin Short Term Repeat (STR) marker results were extracted from Cellosaurus (Bairoch 2018) (3/27/2020). We filtered for human and mouse cell lines (n=109,426 unique accessions). Cell line names and synonyms were converted to lowercase. Where multiple different accessions have the same name (n=31 names, 760 synonyms) or the same names with different punctuation (n=294 names, 579 synonyms), we map every instance to all accessions. If a synonym matches a cell line name it does not share an accession with (n=412 exactly, 91 with different punctuation), we map that name only to the cell line accession where the name belongs, but not to both. A subset of the identically named cell lines have the same parent cell line (n=77 names, 739 synonyms); in this case, the parent cell line is used for the subsequent analysis steps.

We mapped samples to cell lines by matching values to Cellosaurus names and synonyms. We performed mapping using three sets of attributes, of decreasing specificity: (1) attribute pairs with keys containing “cell” and “line”, (2) attribute pairs with values containing “cell” and “line”, and (3) attribute pairs that mention the word “cell”. Human and mouse data were mapped separately, using the appropriate Cellosaurus subset. Prior to analysis, mouse strain names, common stopwords, and cell line names/synonyms that consisted of all numeric characters were removed. We first used exact matches between the value and cell line names (>= 3 characters) for attribute pairs with a cell line key. Then, cell line mentions were detected using n-gram matching (n=1,2,3) between the attribute value (>3 characters) to a cell line or synonym.

##### 3-1-2 Labeling sample source type

Based on the presence of exact lexical matches to key terms in the sample metadata, we automatically assigned samples to one of: tissue, stem cell, xenograft, cancer cell, cell line, primary cell, or other. Cell line data was divided into “named” and “unnamed” cell lines, where named cell lines map to a Cellosaurus identifier (see Supplementary Figure 5 for logic and Supplementary Table 4 for counts by sample type).

#### 3-2 Examining cell line sex

Using our normalized cell line labels, we can compare the reference sex of a cell line from Cellosaurus to the imputed sex from our expression data. For this analysis, we examined cell line samples that were both labeled as a cell line using our expression model and mapped to a Cellosaurus ontology label based on its metadata.

Two types of reference sex labels from Cellosaurus were compared to imputed sex labels from our model; these include *donor* sex, the sex of the donor the cell line was derived from, and *recorded* sex, which is the sex of cell line samples derived from Short Tandem Repeat (STR) profiling of the amelogenin genes (Sullivan et al. 1993). Samples are labeled as “both” if multiple STR analyses have obtained different sex labels.

For cell lines with corresponding data in the Cancer Cell Line Encyclopedia (CCLE) (Barretina et al. 2012), we compared our sample sex scores to CCLE X and Y chromosome copy number (CNV) data (downloaded 9/5/2020).

### 4. Estimation of metadata misannotation

We examined metadata sex label misannotation rates in three ways, described below.

#### 4-1 Comparing mismatch rates in single sex versus mixed sex studies

We examined the rates of sample and study sex label mismatches in large single and mixed sex studies (large is defined as having at least 10 samples). A sample mismatch is a sample with a metadata sex label of male or female and an expression-based sex label above a given threshold (0.7) indicating the opposite sex. A mismatched study means that the study contains at least one mismatched sample. We used a chi-squared test to examine whether the fraction of mismatched samples and studies was significantly different across in mixed versus single sex studies.

#### 4-2 Comparing expression-based methods in mixed sex studies

For large mixed sex studies (at least 5 male and 5 female samples), we compared metadata sex labels with expression labels predicted from the clustering-based methods from Toker et al. (Toker, Feng, and Pavlidis 2016) and massiR (Buckberry et al. 2014) and our own classification method. We conservatively labeled a sample as a mismatch if all of the expression labels disagreed with the metadata label.

#### 4-3 Clustering to identify high confidence swaps in mixed sex studies

While the model predicted probability (“*sample sex score*”) provides an estimate that a sample is a particular sex, we find there is a lot of study-to-study heterogeneity in the distribution of these sample sex scores. By clustering the sample sex scores, we can leverage information about their distribution within a mixed sex study to obtain a probability estimate that a sample belongs to either the male or female cluster (Figure 4). This provides a more local estimate of mis-annotated samples at a study level and allows us to identify high confidence swaps. To obtain a probability estimate that a sample belongs to a cluster, we fit a mixture of one-dimensional Gaussians.

**Figure 1.**
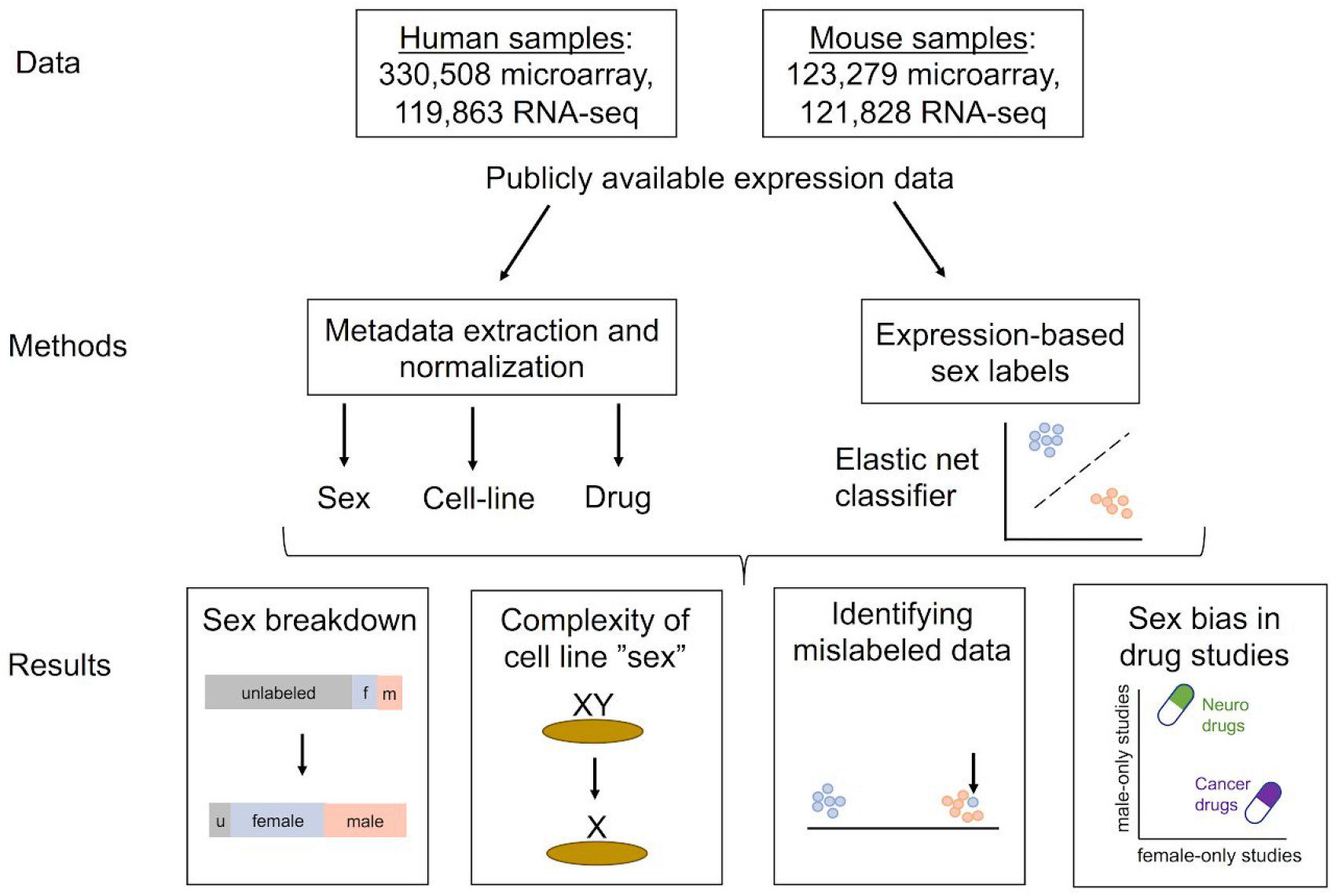
Schematic of study analysis. All human and mouse samples from refine-bio were gathered, metadata extracted and normalized, and sex labels inferred based on expression. This allowed us to examine sex breakdown, cell line sex, and bias in drug studies.

**Figure 2.**
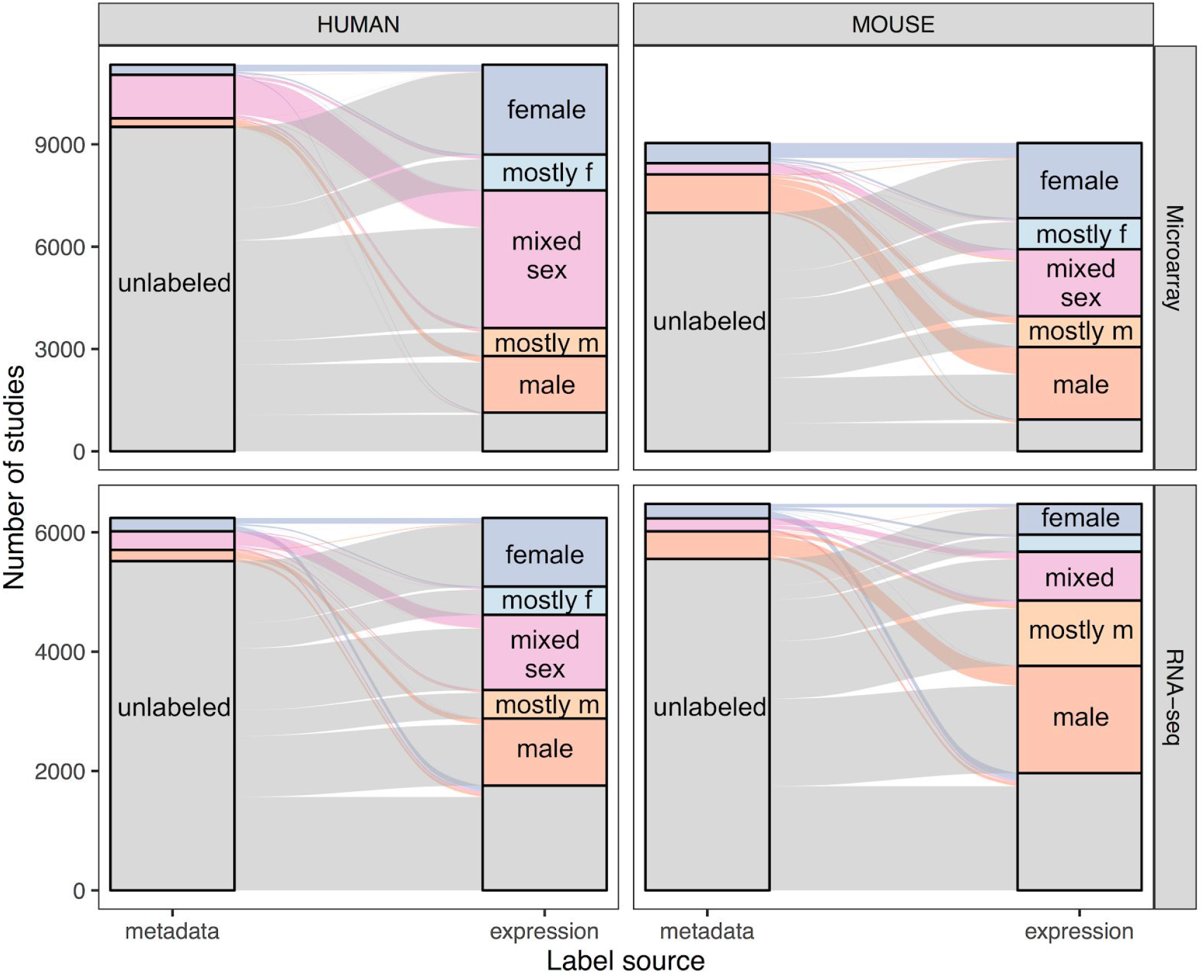
Alluvial diagram showing the breakdown of study sex labels in the metadata (left of each panel) and after expression based labeling (right of each panel). The flow is colored by the initial metadata labels and helps trace whether there is a “change” in labels. For the majority of studies with metadata labels, the labels match the imputed expression labels. The results are shown for both human and mouse (columns) and in microarray and RNA-seq (rows). On the left of each panel is the metadata sex breakdown, on the right is the breakdown after expression labeling. Gray indicates that a study is missing sex labels (for n <=60, more than half of the study labels are missing or unlabeled, for n>60, there are fewer than 30 labels), dark blue means the samples in the study are female-only, dark orange is male-only, and pink is mixed-sex. Mostly male (light orange) and mostly female (light blue) indicate that more than 80% of the samples labeled in that study are of that sex. For the metadata breakdown, the numbers of studies in mostly-male and mostly-female categories was small (8-87 studies) and were grouped into mixed sex for ease of visualization. (For a similar figure with sample sex breakdown see Supplementary Figure 1.)

**Figure 3.**
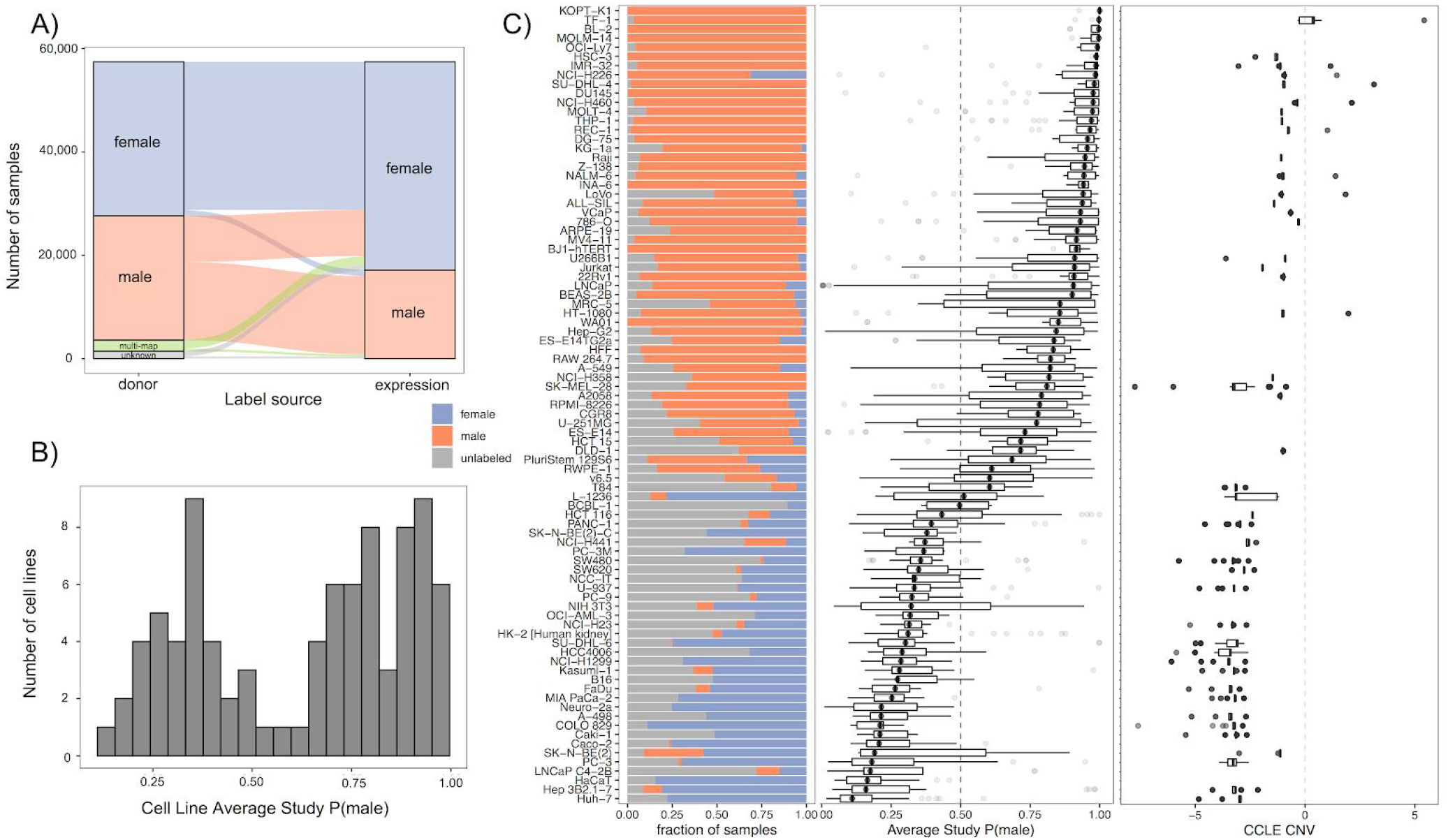
Y chromosome loss is prevalent and variable across cell lines. A) Cell line sex label switching. The sex of the donor cell line is on the left and the imputed sex is on the right. Samples are divided into female (blue) and male (orange). Additionally for donor cell sex labels, we include samples with unknown sex (gray) and samples with metadata mapping to more than one cell line (green). B) Distribution of average study sex scores (e.g. P(male) for a sample) for male cell lines shows a bimodal pattern, indicating that many of these cell lines appear “female-like”. (see Supplementary Table 7 for a list). C) Profiles of Y chromosome loss across male cell lines with more than five studies. Each cell line is a row. From left to right, the first panel shows the fraction of male (orange), female (blue), and unlabeled (gray) samples, the second shows the distribution of study average sample sex scores, and the third shows the cell line specific distributions of Y chromosome gene copy number (CNV) from the Cancer Cell Line Encyclopedia (Barretina et al. 2012). Overall, this demonstrates the highly variable and cell-specific nature of Y chromosome loss.

**Figure 4.**
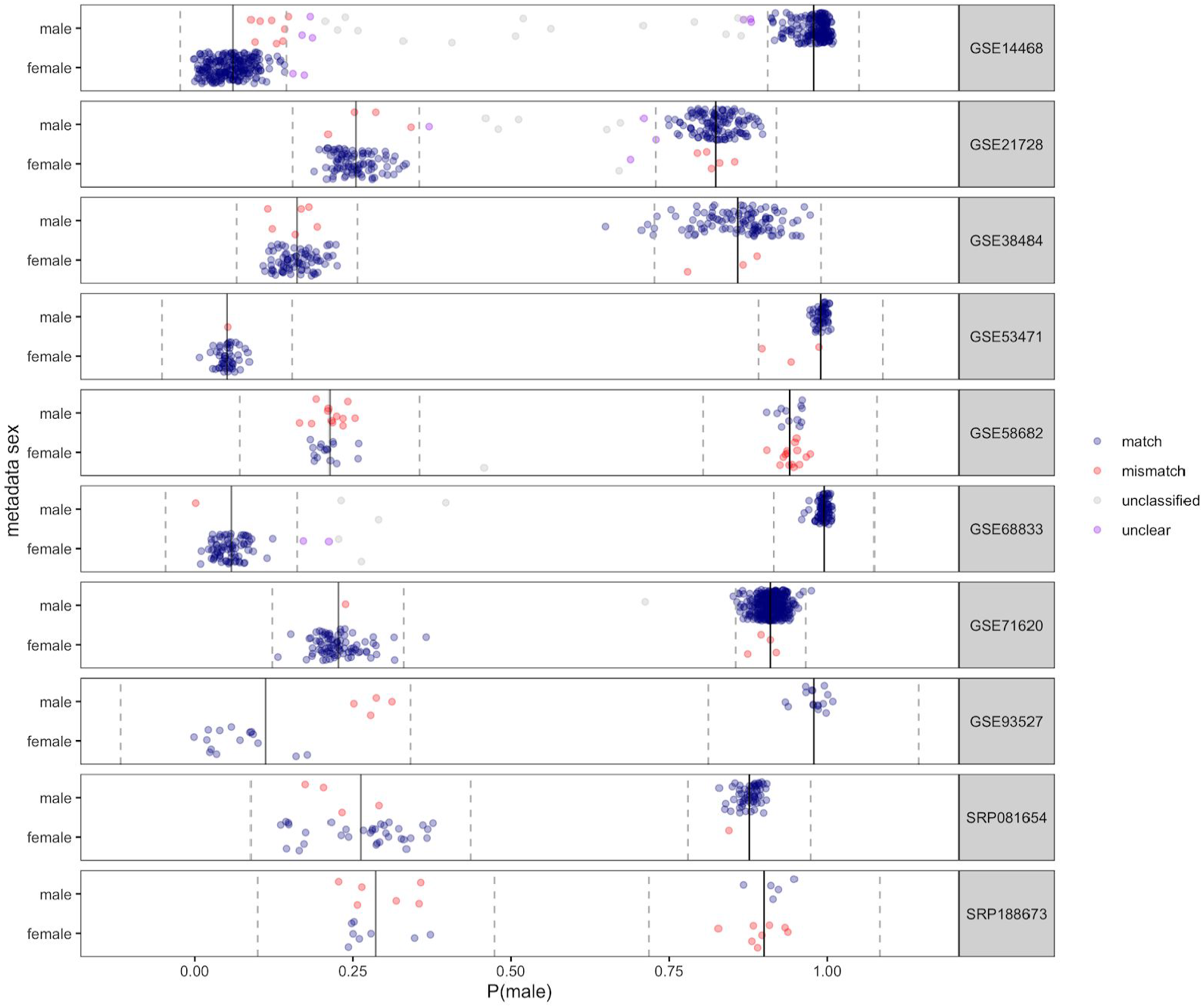
Leveraging within-study distributions of sample sex scores to identify high-confidence mislabeled samples. Each row is a study (randomly sampled from the list of mixed sex tissue studies with multiple clusters). Samples are separated by metadata sex (on the y axis) and our model sample sex score (P(male)) (on the x axis). Samples are colored by whether they show a high confidence (as indicated by a P(sample belongs to cluster) > 0.95) “match” (blue) or “mismatch” (red) between the metadata and expression-based sex; samples that were not classified by the model are labeled “unclassified” (gray), classified samples that do not pass the 0.95 threshold for their cluster are labeled “unclear” (purple). Clustering was obtained by fitting a mixture of Gaussians; and the estimated mean (solid line) and 95% confidence interval (dashed line) for each cluster is shown.

To identify high confidence misannotated samples in mixed sex studies, we used Gaussian model based clustering (R package Mclust (Scrucca et al. 2016)) to cluster the sample sex scores within a study. We performed clustering by fitting a mixture of Gaussians using the unequal variances model. We used the default prior with a larger scale parameter (scale=0.15) to account for the spread of samples. Noise was added in the case that sample sex scores fell in the unclassified category (0.3 < p(male) < 0.7) and was initialized to the set of these samples; however, for studies with >⅓ samples unclassified, we did not include noise terms to help with convergence. The number of Gaussians (n=1 or 2) and the best model for each study was selected using Bayesian Information Criterion (BIC). We additionally filtered studies with little separation between clusters, removing studies where the difference in means between the two clusters’ sample sex scores was less than 0.3 (n=2 studies). For the remaining studies, we set a cutoff posterior probability of 0.95 for assignment of a sample to a cluster, in cases where the metadata sex does not match that cluster, we have a “mismatched” sample. The remaining samples are labeled “unclassified” (if the model estimated that they were noise) or “unclear” (assigned a cluster by the model but with probability < 0.95).

### 5 Sex bias in drug data

#### 5-1 Drug labeling

##### 5-1-1 Study drug mention labeling

Studies were labeled as having a drug mention if the metadata contained a drug name. DrugBank (Wishart et al. 2018) (date accessed: 4/14/2019) XML data was downloaded and synonyms and drug names parsed. Names or synonyms 3 or fewer characters long were discarded, as well as common stop words. Then, we used n-gram matching (n=1,2,3) to map between the metadata text (either study or sample) and a drug or synonym.

To find drug-containing studies with high sensitivity, we labeled studies based on *drug mentions* in their metadata. Study mapping used the text from study title and description fields. Out of 44,184 total studies, 7665 (17.3%) contained a drug mention (1104 drugs).

##### 5-1-2 Sample-level annotation of drug labeling

To build a high specificity dataset, we also performed sample level labeling using the refine.bio harmonized “treatment” and “compound” fields, as well as the sample title.

We created a library of common control terms for sample mapping. This vocabulary consists of the following words:

> “none”, “control”, “untreated”, “dmso”, “na”, “placebo”, “saline”, “pbs”, “mock”, “baseline”, “unstimulated”, “etoh”, “ethanol”, “ctrl”, “non-treated”, “vehicle”, “ctl”, “no treatment”

We then identified the subset of studies where the drug mentioned in a treatment field is the drug mentioned in the study; we call these *drug exposure studies*.

#### 5-2 Assessment of sex bias in drug data

Anatomic Therapeutic Class (ATC) drug mappings were extracted from DrugBank. The enrichment of male only vs female only, and single vs mixed sex studies in each class was assessed separately using chi-squared tests. Prior to running tests, we removed classes with less than 5 samples in a category. We filtered for a corrected chisq p-value < 0.05 using Bonferroni correction on the number of tests (n=48). We also grouped by drug and calculated the fraction of male-only and female-only studies for each drug.

## RESULTS

### 1. Aggregation of publicly available mouse and human expression data

We downloaded expression data and metadata for all human and mouse samples and studies from refine-bio. After filtering (see Methods 1-1), this resulted in 330,508 and 123,279 microarray samples (spanning 11,333 and 9,303 studies) and 119,863 and 121,828 RNA-seq samples (spanning 6,240 and 6,477 studies) respectively (Figure 1).

### 2. Large-scale sex labeling of mouse and human data

#### 2-1 The majority of samples are missing metadata sex labels

We extracted and examined the missing of metadata sex labels in all mouse and human data present in refine.bio (see Methods 2-1). Our analysis indicates that 68-86% percent of samples and 77-88% percent of their corresponding studies are missing sex labels (Figure 2, S1, Tables S1A and B). In the absence of metadata sex labels, we can neither assess inclusion of males and females in studies nor examine whether there are sex-related effects.

#### 2-2 Inferring sample sex from expression data

Existing methods for inferring missing sex labels from expression data are limited to mixed sex studies (Toker, Feng, and Pavlidis 2016; Buckberry et al. 2014) (and as a result cannot be applied *a priori*) or to specific data or platforms (Giles et al. 2017; Ellis et al. 2018). In order to label all publicly available expression data, we trained penalized logistic regression models to impute sample sex from the expression of X and Y chromosome genes. The predicted value from the model corresponds to the model’s predicted probability of that sample being male (P(male) or P(sex=1) using the standard coding where female is 0 and male is 1); we refer to this value as a *sample sex score* and leverage this score to better understand the distributions of samples. In many cases we both use this to label sample sex, assigning samples to male or female at a certain threshold cutoff, and examine the distribution of these scores.

We assessed the accuracy of our model in a randomly held-out test set and in various subsets of the data, compared to all metadata labels (agreement 90.8-93.9%). We additionally looked at the performance for the subsets of these labels in single sex studies (agreement 88.7-96.7%), mixed sex studies (91.5-96.5%), and, in human, manually annotated sex labels from a previous analysis (94.2%) (Giles et al. 2017). As expected, at more stringent cutoffs for assigning sample sex, we achieve higher concordance at the expense of leaving a portion of samples unlabeled (Supplementary Figure 2, values in Supplementary Table 2B). We selected a threshold of 0.7 to correspond with approximately 95% accuracy across the datasets. Our models show good performance in most platforms; however, 6 of 62 platforms (covering <3% of all samples) have very poor performance (accuracy < 70%, see Supplementary Figure 3 and Supplementary Table 3A for platform-specific accuracy). We filtered to remove these “problem platforms”, and had 92-97% accuracy with 70-71% of RNA-seq and 91-93% of microarray data labeled at a model threshold of 0.7 (Supplementary Table 3B).

#### 2-3 Sex labeling mixed sex or pooled samples

For a small fraction of samples (0.05-4.2%), their metadata indicates that they are pooled or mixed sex samples; this practice is more common in mouse data, as samples are often pooled from mice of the same strain before analysis, and it is used to both increase signal and reduce the number of expression samples (and thereby the cost). Pooling often includes samples from both sexes, but this is highly variable. We sought to examine our models’ predictions on these mixed or pooled samples. The models’ sample sex score distributions are significantly different in mixed/pooled samples versus male or female labeled samples; where the model distribution appears almost uniform for mouse pooled samples (see Supplementary Figure 4). At a threshold of 0.7, pooled samples fall into the unlabeled category at higher rates than samples with male and female metadata sex (in mouse microarray data, 28.8% of pooled versus 5.4 and 7.7% of female and male samples are not labeled).

#### 2-4 Sex breakdown shows slight female bias in humans and male bias in mice

We applied these models to all publicly available human and mouse expression data and found that the overall sex breakdown is slightly female-biased (52.1%) in humans and male-biased (62.5%) in mice (Figure S1, Table S1A). At a study-level, in humans, the majority of studies are mixed sex (41.0% mixed sex vs 33.3% female-only and 18.4% male-only), while in mice, studies are evenly split with about one third in each category (31.0% female-only, 30.4% male-only, 29.6% mixed sex). This pattern does not appear to change over time (Supplementary Figure 5).

### 3. Cell line “sex” is complex

We performed named entity recognition to identify cell lines based on the metadata and mapped 74,140 out of 99,426 human samples and 8,433 out of 17,061 mouse samples to Cellosaurus identifiers. For human RNA-seq samples, this showed high concordance with MetaSRA (76.5% exact matching, 14,390 out of 18,821 samples).

#### 3-1 Our analysis supports previous observations that cell line “sex” is fundamentally different from tissue sex

Previous studies (Shah, McCormack, and Bradbury 2014) have encouraged researchers to report the sex of their cell lines; however, several other studies have shown that cell line sex can be a complicated concept, in part because cell lines often lose their Y chromosomes in culture (Molaro and Malik 2017). We find that many samples with reference sex male are labeled as female based on expression data (37.8%) while relatively few samples have reference sex female and are labeled male (4.24%, Figure 3A, Supplementary Figure 8). This pattern of cell line “switching” from male to female matches patterns that have already been shown on a smaller scale; namely, many cell lines lose Y chromosomes.

Overall, we see significant enrichment of the overall proportion of female imputed labels in cell line versus tissue samples (Supplementary Figure 7A, p<0.05 for human and mouse microarray and human RNA-seq, N.S. for mouse RNA-seq). Additionally, the distributions of sample sex scores for cell line vs tissue shows increased female scores and a wider distribution of scores in cell line versus tissue data (Supplementary Figure 7B).

#### 3-2. Our analysis provides evidence of cell-line specific patterns of Y chromosome loss

Across male cell lines, the fraction of inferred male versus inferred female samples varies greatly, with certain cell lines appearing more “male-like” (e.g., THP-1, OCI-Ly7), “female-like” (e.g. KYSE-30, HaCaT), or cell lines appearing to belong more in the middle with samples belonging to both (A549, HCT-116) (Figure 3B-C). We further examined 87 cell lines from male donors that had a large number of studies and samples (at least five studies, with at least 3 samples in each study) (Supplementary Table 5). Of these cell lines, 45 are included in Cancer Cell Line Encyclopedia (Barretina et al. 2012). On a cell line level, our sample sex score predictions correlate with CCLE Y but not X chromosome copy number (Spearman correlation of medians for Y: 0.774, p< 4.58 x 10^-10^, and X: −0.0944, p = 0.537) (Figure 3C).

### 4. Our models allow for improved detection of mis-annotated data

We can estimate metadata mis-annotation rates at scale by comparing our inferred sex labels to metadata sex labels. First, we examined the mismatch rates in mixed sex versus single sex tissue studies and found significantly higher mismatch rates in samples from human mixed sex studies (4.87%) than single sex studies (2.03%) (Chi-squared p-value < 1.69 *10^-48). By contrast, in mouse studies, mismatch rates for samples in single sex studies (3.42%) exceeded that of mixed sex (2.58%) studies (p < 9.70*10^-6) (Supplementary Table 6A).

Second, we created a dataset of large mixed sex studies with metadata labels, consisting of 6066 mouse and 8658 human samples (168 and 163 studies respectively), which we sex labeled with the Toker and massiR methods, as well as our own (see Methods 4-2). Across these samples, 4.74% of human and 6.64% of mouse samples had metadata sex labels that did not match any of the predicted expression-based labels. At a study level, 35.0% of human and 16.7% of mouse studies contained at least one mismatched sample (Supplementary Table 6B).

Third, to label estimate the probability that an individual sample is mislabeled in a mixed sex study, we fit a mixture of Gaussians to each study-level distribution of sample sex scores (see Methods 4-3). This allows us to identify high confidence mismatches at a particular probability threshold (>0.95) by examining the concordance between metadata labels and expression labels. After discarding mixed sex studies where the best model was a single Gaussian (409 or 26.9% of mixed sex studies -- by comparison, we also clustered single sex studies and found 94.0% of single sex studies were best modeled by a single Gaussian), we estimate that 2.04% of mouse and 3.06% of human samples are mislabeled in these studies. At a study level, this corresponds to 26.0% of mouse and 51.7% of human studies containing at least one high confidence mismatched sample (Supplementary Table 6C).

### 5. Sex breakdown of drugs in tissue data

#### 5-1 Labeling drug studies using metadata

We identified all gene expression studies containing drug mentions by applying named entity recognition and normalization to the study and sample metadata (see Methods 5-1). We labeled studies with *drug mentions* (n=7,665), which contain a drug in the study abstract or title, and *drug exposure* studies (n=1,095) which include a drug in the sample treatment field.

Out of the 95,788 human and mouse samples with non-null treatment fields (spanning 9,130 unique treatment labels), 30,000 mapped to only control terms (1,144 unique) and 463 mapped to DrugBank drugs and control terms (91 unique), 12,692 mapped to just drugs (1,410 unique treatment labels, mapping to 417 drugs), leaving 52,633 unmapped treatment labels (6,485 unique). 96 additional samples mapped using compound or title.

The overlap between the studies with drug mentions (n=7,665) and treatment studies (n=1,095) is 757 studies; 557 of these map to the same drug or drug(s) and 159 map to at least one of the same drugs. After filtering for only overlapping drugs, there are 844 unique study-drug pairs spanning 319 drugs and 715 studies. We include a list of these *drug exposure studies* in Supplementary Table 6.

#### 5-2 The sex breakdown of drug studies shows limited sex bias overall

We then asked the question of whether global patterns of sex breakdown continue in the context of drug studies. We found that this was the case; with slightly increased female-bias in human studies (52.7% overall, 59.9% in drug studies) and male-bias in mouse studies (52.0% overall, 57.1% of drug studies) (Supplementary Figure 9). This pattern varies across drug Anatomic Therapeutic Class (ATC), with female-only studies enriched in human and mouse genitourinary system drugs (class G, p < 10^-8^ and p < 10^-16^ respectively) and in human cancer/immune system drugs (L, p < 10^-7^), and male-only studies were enriched in mouse nervous system drugs (N, p < 10^-7^) (Figure 5). These patterns match what is seen in drug exposure studies (Supplementary Figure 10A,C) and a previous dataset of manually curated drug expression studies ((Wang et al. 2016), n=472) (Supplementary Figure 10B,D).

**Figure 5.**
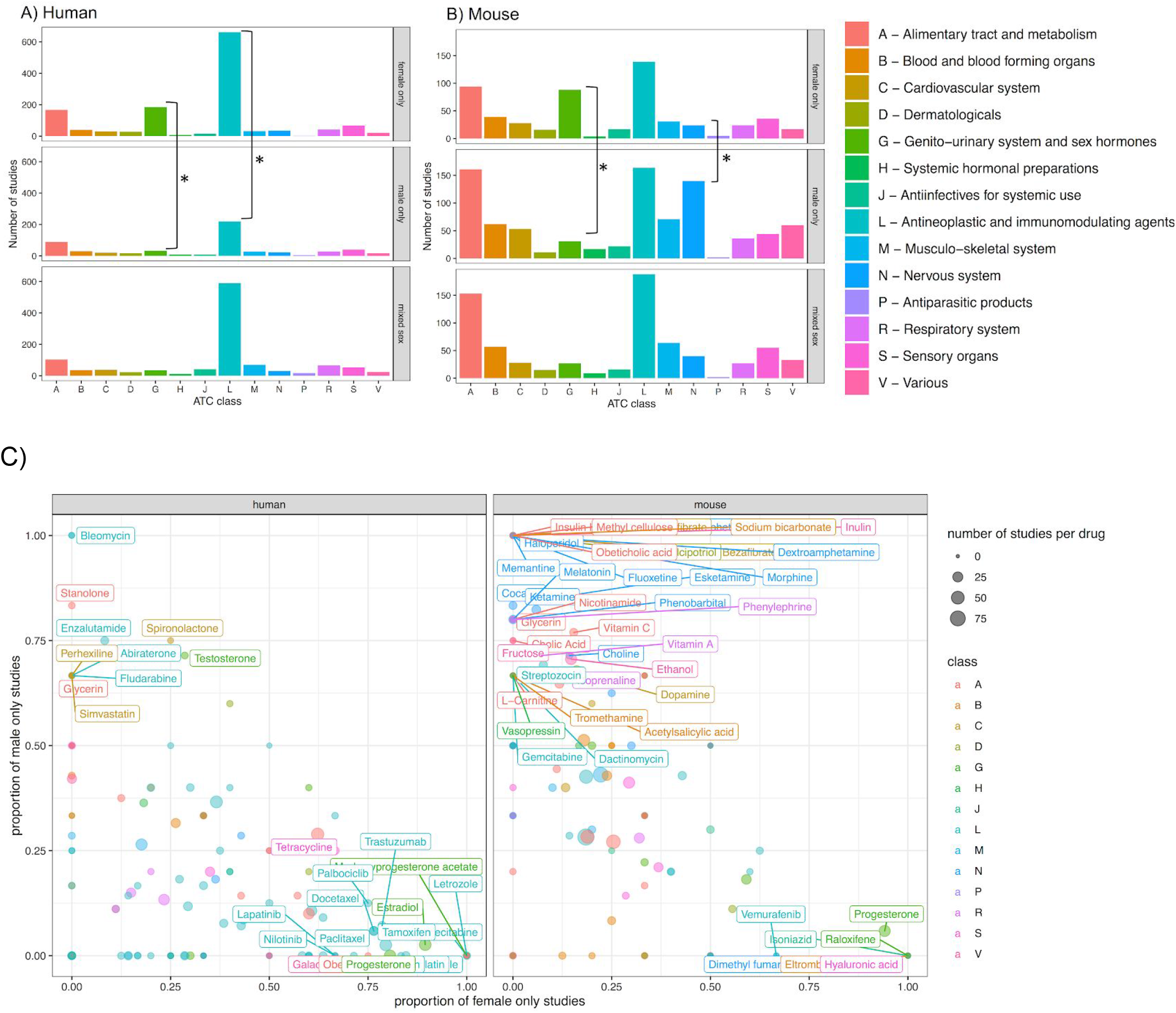
Sex breakdown by drug ATC class in human and mouse data shows enrichment of female only studies in human cancer drugs and male only studies in mouse drugs. Count breakdown of female only, male only, and mixed sex studies by class in humans (A) and mice (B) (* indicates p<0.05 after Bonferroni correction). The fraction of studies for each individual drug is also shown in (C), where the x axis is the fraction of female only studies and the y axis is the fraction of male only studies. Each point is a drug, the size of the point indicates the number of studies that include that drug, and the color of the point is ATC class. Drug name labels are included for drugs with at least three studies and a strongly sex-biased ratio (>⅔ of studies only one sex, <⅓ the other); other labels are omitted for ease of visualization. For a full list of drugs see Supplementary Table 7.

#### 5-3 Human cancer drug studies are female-biased

There are 370 female-only versus 105 male-only studies with cancer drug mentions; however, the majority of cancer drug studies are still mixed sex (n=396). These studies span 143 different drugs and despite being highly female-biased at the study level, 115 of these drugs (80.4%) have at least one mixed sex study.

To determine the possible association between cancer and sex, we examined whether drugs targeting strongly sex-biased cancers (e.g. breast, ovarian, prostate) were associated with more single sex studies. In humans, of the drugs with >70% female-only studies 9 out of 21 are breast cancer specific and 19 out of 21 drugs contain indications for breast or ovarian cancer (Supplementary Table 8A). For drugs with >70% male only studies, in humans 2 out of 8 of these are used for prostate cancer (Enzalutamide, Abiraterone); the remainder are used for multiple types of cancer (this includes Bleomycin which is used for multiple cancers including ovarian).

#### 5-4 Mouse nervous system drug studies are male-biased

In the case of mouse nervous system drugs, we see high male-bias in the proportion of male mice used as study subjects. This pattern of using only male rodents in neuroscience has been previously reported ((Beery and Zucker 2011). At an individual drug level, while the majority drugs either single and mixed sex (n=147, 38.7% of human; n=95, 34.5% of mouse) or mixed sex only studies (n=115, 30.3% of human; n=38, 13.8% of mouse), some drugs showed very high male-bias (10.3% of human and 20.7% of mouse). In nervous system drugs in particular (n=27 in humans, n=37 in mice), this fraction is markedly increased in mouse nervous system drugs, with 13 drugs that are only studied in males versus 5 only in females (see Supplementary Table 8B for a list of neuro drugs). The 13 drugs with male-only studies in mice, two have mixed sex studies in humans (Herion and Carbamazepine), two only have male-only studies in humans (Fluoxetine and Haloperidol), and the remainder have no human studies (Acepromazine, Amphetamine, Levadopa, Memantine, Modafinil, Olanzapine, Quetiapine, Salsalate, Venlafaxine). Fluoxetine is particularly interesting because it is a commonly prescribed antidepressant and is present in eight mouse studies, all of which are male-only. The eight studies are all treatment (fluoxetine) versus control comparisons and collectively contain 128 samples all from mouse brain tissue. In humans, there is only one fluoxetine study and it is also all male.

## DISCUSSION

We inferred sample sex labels at scale from expression data; to our knowledge, our analysis represents the first effort to label all human and mouse microarray and RNA-seq data. We leverage these labels to assess sex bias overall and in drug studies, examine cell line sex, and estimate misannotation rates. Below, we discuss our findings in the context of previous results, and potential limitations of our methodology.

### Labeling expression data at scale

Improving on previous methods, which have high accuracy but are focused on specific data or platforms (Giles et al. 2017; Ellis et al. 2018) or mixed sex studies (Buckberry et al. 2014; Toker, Feng, and Pavlidis 2016), our labeling method has consistent accuracy across the majority of platforms and in both mixed and single sex studies. We leverage the sample and study metadata to label sample sources (e.g. tissue, cell line, primary cell, etc) and map samples and studies to cell line and drug identifiers from Cellosaurus (Bairoch 2018) and DrugBank (Wishart et al. 2006) respectively. Our combined metadata mappings and imputed sex labels allow us to examine the sex breakdown across sample source type, cell lines, and drug-related expression studies.

We found that the overall sex breakdown shows slight female-bias in human samples and male-bias in mouse samples. This breakdown continues at a study level, where the majority of studies are mixed sex but there is still slight female- and male-bias in human and mouse studies respectively. This low overall sex bias is in contrast to what we expected, given the history of clinical trials and mouse studies excluding women, but this is consistent with a previous finding in human RNA-seq data (Ellis et al. 2018). It is important to note our analysis focuses on publicly available expression studies and does not extend to private data from drug company data or clinical trials.

When we examined labels over a fifteen-year period (2004-2019), we found that the missingness of sex labels and patterns of sex bias did not appear to change over time. This was also unexpected because in 2016 there was an NIH mandate to include sex as a variable in preclinical studies and overall there has been increased awareness of its importance. Specific fields have documented improvements in the inclusion of females and reporting of sex (Woitowich, Beery, and Woodruff 2020); it is possible that future analysis grouping by field may show changes in the sex breakdown over time despite the overall pattern seeming consistent.

### Sex labeling cell lines shows Y chromosome loss

In 2014, Shah et al. (Shah, McCormack, and Bradbury 2014) encouraged researchers to include and examine the sex of the cells they are using. Previous studies have shown that certain types of male and female cells in culture respond differently to drugs (Mennecozzi et al. 2015), underscoring the importance of this consideration. However, there is an important distinction between examining the sex of primary or stem cells and examining the sex of transformed cell lines. While the sex of the cell line donor provides some information, cell lines in culture often undergo chromosomal loss or duplication (Xu et al. 2017); in particular loss of Y chromosomes is common (Molaro and Malik 2017)

While the gold standard for cell line sex labeling is PCR-based authentication (Sullivan et al. 1993), in the absence of this information, we can use our models to infer sample sex and identify Y chromosome loss from expression data, allowing for re-examination of existing studies. We found both highly prevalent (43.4% of male cell lines contain at least one female-appearing sample) and variable patterns of Y chromosome loss across cell lines. It is unclear whether this variability across cell lines is due to differences in cell stability (Fasterius and Al-Khalili Szigyarto 2018), passage number (Y chromosome loss is more common after longer time in vitro (Xu et al. 2017)), contamination, or mislabeling. We additionally found that the distribution of sample sex scores in cell lines varies greatly from that of tissues and primary cells, with human primary cell line distributions appearing to more closely resemble tissue than cell lines. Many mouse primary samples had ambiguous sample sex scores, which is likely due to the practice of using pooled samples for mouse primary cultures. Altogether these results highlight the lack of a cell line sex binary and match previous recommendations against using cell lines for examining sex-related effects (Ritz 2017). We performed the remainder of the analysis on cell lines and tissues separately.

While we infer potential Y chromosome loss in cell lines with donor sex male that appear female in our expression data, it is also possible that these cell lines are contaminated, which is very common (Capes-Davis et al. 2010). Proper cell line authentication (via PCR) is necessary to confirm contamination; however, this is not possible during re-analysis of existing studies. Our estimates are limited in that they also do not provide direct copy number information, which is important, as cell lines also often undergo X chromosome duplication or changes in autosomal ploidy (Xu et al. 2017). However, the sex breakdown of individual cell lines correlate with the cell’s Y chromosome reference copy number variation (CNV) statistics from Cancer Cell Line Encyclopedia (CCLE), validating our inferred labels and providing further evidence of cell line specificity.

Another potential factor that may influence sex-related behavor in cell lines is the presence of media, which we did not examine in this analysis. Many medias contain phenol red as an indicator, which has estrogenic properties, and serum, which mimics a pregnancy-like environment, and these may impact the study of sex-related effects (De Souza Santos et al. 2018; Ritz 2017). Examining the effects of media is very challenging as there is great heterogeneity and media types are often underreported or missing specific information (De Souza Santos et al. 2018).

### Estimation of misannotation rates

Metadata mislabeling is a widespread problem. Previous studies (Lohr et al. 2015; Toker, Feng, and Pavlidis 2016) have used imputed expression labels to estimate mislabeling rates. Toker et al found that 46% of the 70 mixed sex datasets they examined contained mislabeled samples, with an overall sample mislabeling rate of 2% (4160 samples). Using three methods, we estimate that 2-5% of samples contain misannotated sex labels and 30-50% of studies contain at least one misannotated sample. Our estimates match previous studies (Toker, Feng, and Pavlidis 2016; Lohr et al. 2015), but expand this analysis to include many more microarray platforms, RNA-seq studies, and mouse samples.

We initially examined the difference in mismatch rates between mixed and single sex tissue studies, with the expectation that mixed sex studies would have higher mismatch rates than single sex studies because of the potential for swapped annotations. This was the case in human but not mouse data. It is unclear why single sex studies show higher mismatch rates in mice and it is possible that this may be a limitation of our metadata sample source identification pipeline. Our approach may have poorer performance in mouse data than human data at discriminating identifying cell line samples (the latter we evaluated through comparison to MetaSRA). Inclusion of cell line samples within the tissue analysis would increase mislabeling rates. Additionally, this difference in rates may be related to the collection of samples from mouse pups, which are hard to sex (Deeney, Powers, and Crombleholme 2016). This raises questions about the proportion of mouse studies that are truly single sex, which requires further investigation, and underscores the importance of using proper authentication for sex determination.

### Including sex as a variable in drug-related studies

The sex breakdown of drug data shows limited sex bias and matches the overall breakdown of samples and studies; however, studies of specific drug classes show sex bias -- particularly mouse nervous system drugs (male-biased) and human cancer drugs (female-biased). While we focused our analysis on studies that mention a drug, which may have low specificity, we found a similar sex breakdown for the subset of these studies with drug information in a treatment field, and a crowd-sourced set of drug studies from CREEDS et al. (Wang et al. 2016). Future work may involve better annotation of drug exposure studies from metadata using deep learning based methods.

While it is of interest to examine whether particular drugs lead to differential gene expression responses between males and females, we are underpowered to identify interaction effects in most of these studies using classical methods because of sample size. However, it is possible that non-parametric methods or pathway and gene set based techniques could be used to examine these effects (Zhou and Wong 2011). Additionally, it is still important to include both sexes in studies regardless of our ability to examine interaction effects (Klein et al. 2015). Our inferred sex labels allow for assessment of these practices. In addition, we hope that these labels can lead to improved selection of studies and inclusion of sex as a variable in re-analysis of public data.

### Analysis Limitations

In development of our sex labeling model, we chose to focus on a model that would allow us to infer sample sex at high accuracy across a wide range of samples, studies, and platforms. Our model has better performance on microarray data, this may be due to the fact that we used similar models across microarray and RNA-seq data by normalizing the count data; however, it is possible that alternate models for count data and normalization methods may perform better, and future work will assess this.

As is common, our model results in one probability score for each sample, allowing for accurate probability cutoffs. However, it is possible that having two predicted scores (for male-like and female-like behavior) could allow for better understanding of sex in cell lines, pooled samples, and samples from intersex individuals. Our analysis uses the expression of X and Y chromosome genes to label sample sex, we did not consider autosomal expression because previous studies have indicated that sex differences in this are generally tissue-specific (Gershoni and Pietrokovski 2017). However, sex differences in autosomal expression, such as determined through methods such as ISEXs (Bongen et al. 2019), is also important for biological understanding and future work may involve including autosomal genes.

As part of our analysis, we also leveraged metadata to map samples and studies to sample source types, cell lines, and drugs. For mappings of samples to cell line and drug labels, we required an exact match with a cell line name or synonym, and for sample source type, we also required exact lexical matches for many of the sample source categories, resulting in many unlabeled or “other” samples. Use of state-of-the-art biomedical named entity recognition and normalization methods may help to improve the accuracy and sensitivity of these labels.

We focused on sex bias in publicly available human and mouse expression data in refine.bio because we wanted to examine sex bias in data from both humans and a mammalian model organism. Mice have the largest amount of available expression data and are often used in drug development. We labeled a large fraction of these data; however, a small subset of samples could not be labeled due platform challenges or missing or poor quality expression data. Since this is a small fraction, and missing data is spread across many studies, we do not expect that labeling these data will change our conclusions; despite this, continued efforts will attempt to “rescue” these missing data. While we found only slight overall bias in publicly available mouse and human expression data, it is possible that public data for other organisms or proprietary data show different patterns of bias. In the future, we hope to extend our methods to sex label publicly available samples from other organisms with XY determination systems in order to further aid in study re-analysis and assessment of sex bias.

## CODE & DATA AVAILABILITY

The sample sex, source, drug, and cell line labels will be made publicly available through refine.bio (https://www.refine.bio/) to allow researchers to examine these labels and use them to better search for and re-analyze existing studies. The code used for labeling and analysis is publicly available at https://github.com/erflynn/sl_label. Cell line annotations of potential Y chromosome loss and drug-study mappings are included in the supplement.

## ACKNOWLEDGEMENTS

E.F. is supported by NIH NLM F31 Fellowship F31LM013053 and the Stanford Data Science program. A.C. was partially supported by the Stanford Bio-X Undergraduate Summer Research Program. R.B.A is supported by the NIH GM 102365, LM 005652 and the Chan Zuckerberg Biohub. The majority of the computing for this project was performed on the Stanford University Sherlock cluster. We would like to thank the Stanford Research Computing Center for providing the computational resources that contributed to these research results. We would also like to thank Dr. Casey Greene and the refine.bio resource for help with this project.

## AUTHOR CONTRIBUTIONS

E.F. and A.C. conceived of the study together. E.F. performed metadata normalization, trained and applied the models, and performed some of the follow up analysis. A.C. helped with assessing the method and performing downstream analysis. R.B.A. supervised the project. All authors contributed to writing the manuscript.

## REFERENCES

Bairoch, Amos. 2018. “The Cellosaurus, a Cell-Line Knowledge Resource.” Journal of Biomolecular Techniques: JBT 29 (2): 25–38.

Barretina, Jordi, Giordano Caponigro, Nicolas Stransky, Kavitha Venkatesan, Adam A. Margolin, Sungjoon Kim, Christopher J. Wilson, et al. 2012. “The Cancer Cell Line Encyclopedia Enables Predictive Modelling of Anticancer Drug Sensitivity.” Nature 483 (7391): 603–7.

Beery, Annaliese K., and Irving Zucker. 2011. “Sex Bias in Neuroscience and Biomedical Research.” Neuroscience and Biobehavioral Reviews 35 (3): 565–72.

Bernstein, Matthew N., Anhai Doan, and Colin N. Dewey. 2017. “MetaSRA: Normalized Human Sample-Specific Metadata for the Sequence Read Archive.” Bioinformatics 33 (18): 2914–23.

Bongen, Erika, Haley Lucian, Avani Khatri, Gabriela K. Fragiadakis, Zachary B. Bjornson, Garry P. Nolan, Paul J. Utz, and Purvesh Khatri. 2019. “Sex Differences in the Blood Transcriptome Identify Robust Changes in Immune Cell Proportions with Aging and Influenza Infection.” Cell Reports 29 (7): 1961–73.e4.

Brazma, Alvis, Pascal Hingamp, John Quackenbush, Gavin Sherlock, Paul Spellman, Chris Stoeckert, John Aach, et al. 2001. “Minimum Information about a Microarray Experiment (MIAME)—toward Standards for Microarray Data.” Nature Genetics 29 (4): 365–71.

Brazma, Alvis, Helen Parkinson, Ugis Sarkans, Mohammadreza Shojatalab, Jaak Vilo, Niran Abeygunawardena, Ele Holloway, et al. 2003. “ArrayExpress—a Public Repository for Microarray Gene Expression Data at the EBI.” Nucleic Acids Research 31 (1): 68–71.

Buckberry, Sam, Stephen J. Bent, Tina Bianco-Miotto, and Claire T. Roberts. 2014. “massiR: Array Datasets.” http://www.academia.edu/download/41619451/massiR_a_method_for_predicting_the_sex_o20160127-31079-18mcqr1.pdf.

Capes-Davis, Amanda, George Theodosopoulos, Isobel Atkin, Hans G. Drexler, Arihiro Kohara, Roderick A. F. MacLeod, John R. Masters, et al. 2010. “Check Your Cultures! A List of Cross-Contaminated or Misidentified Cell Lines.” International Journal of Cancer 127 (1): 1–8.

Chengalvala, Murty V., Vargheese M. Chennathukuzhi, Daniel S. Johnston, Panayiotis E. Stevis, and Gregory S. Kopf. 2007. “Gene Expression Profiling and Its Practice in Drug Development.” Current Genomics 8 (4): 262–70.

Clayton, Janine A., and Francis S. Collins. 2014. “Policy: NIH to Balance Sex in Cell and Animal Studies.” Nature 509 (7500): 282–83.

Collado-Torres, Leonardo, Abhinav Nellore, Kai Kammers, Shannon E. Ellis, Margaret A. Taub, Kasper D. Hansen, Andrew E. Jaffe, Ben Langmead, and Jeffrey T. Leek. 2017. “Reproducible RNA-Seq Analysis Using recount2.” Nature Biotechnology 35 (4): 319–21.

Deeney, Scott, Kyle N. Powers, and Timothy M. Crombleholme. 2016. “A Comparison of Sexing Methods in Fetal Mice.” Lab Animal 45 (10): 380–84.

De Souza Santos, Roberta, Aaron P. Frank, Biff F. Palmer, and Deborah J. Clegg. 2018. “Sex and Media: Considerations for Cell Culture Studies.” ALTEX 35 (4): 435–40.

Edgar, Ron, Michael Domrachev, and Alex E. Lash. 2002. “Gene Expression Omnibus: NCBI Gene Expression and Hybridization Array Data Repository.” Nucleic Acids Research 30 (1): 207–10.

Ellis, Shannon E., Leonardo Collado-Torres, Andrew Jaffe, and Jeffrey T. Leek. 2018. “Improving the Value of Public RNA-Seq Expression Data by Phenotype Prediction.” Nucleic Acids Research 46 (9): e54.

Enache, Oana M., David L. Lahr, Ted E. Natoli, Lev Litichevskiy, David Wadden, Corey Flynn, Joshua Gould, Jacob K. Asiedu, Rajiv Narayan, and Aravind Subramanian. 2019. “The GCTx Format and cmap{Py, R, M, J} Packages: Resources for Optimized Storage and Integrated Traversal of Annotated Dense Matrices.” Bioinformatics 35 (8): 1427–29.

Fasterius, Erik, and Cristina Al-Khalili Szigyarto. 2018. “Analysis of Public RNA-Sequencing Data Reveals Biological Consequences of Genetic Heterogeneity in Cell Line Populations.” Scientific Reports 8 (1): 11226.

Feldman, Sergey, Waleed Ammar, Kyle Lo, Elly Trepman, Madeleine van Zuylen, and Oren Etzioni. 2019. “Quantifying Sex Bias in Clinical Studies at Scale With Automated Data Extraction.” JAMA Network Open 2 (7): e196700.

Friedman, Jerome H., T. J. Hastie, and R. J. Tibshirani. 2010. “Glmnet: Lasso and Elastic-Net Regularized Generalized Linear Models, 2010b.” URL http://CRAN.R-Project.Org/package=Glmnet. R Package Version, 1–1.

Gershoni, Moran, and Shmuel Pietrokovski. 2017. “The Landscape of Sex-Differential Transcriptome and Its Consequent Selection in Human Adults.” BMC Biology 15 (1): 7.

Giles, Cory B., Chase A. Brown, Michael Ripperger, Zane Dennis, Xiavan Roopnarinesingh, Hunter Porter, Aleksandra Perz, and Jonathan D. Wren. 2017. “ALE: Automated Label Extraction from GEO Metadata.” BMC Bioinformatics 18 (Suppl 14): 509.

Greene, Casey S., Dongbo Hu, Richard W. W. Jones, Stephanie Liu, David S. Mejia, Rob Patro, Stephen R. Piccolo, Ariel Rodriguez Romero, Hirak Sarkar, Candace L. Savonen, Jaclyn N. Taroni, William E. Vauclain, Deepashree Venkatesh Prasad, Kurt G. Wheeler. refine.bio: a resource of uniformly processed publicly available gene expression datasets. https://www.refine.bio

Kim, Alison M., Candace M. Tingen, and Teresa K. Woodruff. 2010. “Sex Bias in Trials and Treatment Must End.” Nature 465 (7299): 688–89.

Klein, Sabra L., Londa Schiebinger, Marcia L. Stefanick, Larry Cahill, Jayne Danska, Geert J. de Vries, Melina R. Kibbe, et al. 2015. “Opinion: Sex Inclusion in Basic Research Drives Discovery.” Proceedings of the National Academy of Sciences of the United States of America 112 (17): 5257–58.

Kong, Betty Y., Isabel M. Haugh, Bethanee J. Schlosser, Spiro Getsios, and Amy S. Paller. 2016. “Mind the Gap: Sex Bias in Basic Skin Research.” The Journal of Investigative Dermatology 136 (1): 12–14.

Lachmann, Alexander, Denis Torre, Alexandra B. Keenan, Kathleen M. Jagodnik, Hoyjin J. Lee, Lily Wang, Moshe C. Silverstein, and Avi Ma’ayan. 2018. “Massive Mining of Publicly Available RNA-Seq Data from Human and Mouse.” Nature Communications 9 (1): 1366.

Leinonen, Rasko, Ruth Akhtar, Ewan Birney, Lawrence Bower, Ana Cerdeno-Tárraga, Ying Cheng, Iain Cleland, et al. 2011. “The European Nucleotide Archive.” Nucleic Acids Research 39 (Database issue): D28–31.

Leinonen, Rasko, Hideaki Sugawara, Martin Shumway, and International Nucleotide Sequence Database Collaboration. 2010. “The Sequence Read Archive.” Nucleic Acids Research 39 (suppl_1): D19–21.

Lohr, Miriam, Birte Hellwig, Karolina Edlund, Johanna S. M. Mattsson, Johan Botling, Marcus Schmidt, Jan G. Hengstler, Patrick Micke, and Jörg Rahnenführer. 2015. “Identification of Sample Annotation Errors in Gene Expression Datasets.” Archives of Toxicology 89 (12): 2265–72.

Mamlouk, Gabriella M., David M. Dorris, Lily R. Barrett, and John Meitzen. 2020. “Sex Bias and Omission in Neuroscience Research Is Influenced by Research Model and Journal, but Not Reported NIH Funding.” Frontiers in Neuroendocrinology 57 (April): 100835.

Mennecozzi, Milena, Brigitte Landesmann, Taina Palosaari, Georgina Harris, and Maurice Whelan. 2015. “Sex Differences in Liver Toxicity—Do Female and Male Human Primary Hepatocytes React Differently to Toxicants In Vitro?” PloS One 10 (4): e0122786.

Mogil, Jeffrey S., and Mona Lisa Chanda. 2005. “The Case for the Inclusion of Female Subjects in Basic Science Studies of Pain.” Pain 117 (1-2): 1–5.

Molaro, Antoine, and Harmit S. Malik. 2017. “Culture Shock.” eLife. https://doi.org/10.7554/eLife.33312.

Patro, Rob, Geet Duggal, Michael I. Love, Rafael A. Irizarry, and Carl Kingsford. 2017. “Salmon Provides Fast and Bias-Aware Quantification of Transcript Expression.” Nature Methods 14 (4): 417–19.

Perry, P. O. 2009. “Bcv: Cross-Validation for the SVD (Bi-Cross-Validation).” R package version.

Peterson, Ryan A., and Joseph E. Cavanaugh. 2019. “Ordered Quantile Normalization: A Semiparametric Transformation Built for the Cross-Validation Era.” Journal of Applied Statistics, June, 1–16.

Prakash, Vivek S., Neel A. Mansukhani, Irene B. Helenowski, Teresa K. Woodruff, and Melina R. Kibbe. 2018. “Sex Bias in Interventional Clinical Trials.” Journal of Women’s Health 27 (11): 1342–48.

Ramasamy, Adaikalavan, Adrian Mondry, Chris C. Holmes, and Douglas G. Altman. 2008. “Key Issues in Conducting a Meta-Analysis of Gene Expression Microarray Datasets.” PLoS Medicine 5 (9): e184.

Ritz, Stacey A. 2017. “Complexities of Addressing Sex in Cell Culture Research.” Signs: Journal of Women in Culture and Society 42 (2): 307–27.

Roberts, S., and G. Nowak. 2014. “Stabilizing the Lasso against Cross-Validation Variability.” Computational Statistics & Data Analysis 70 (February): 198–211.

Scrucca, Luca, Michael Fop, T. Brendan Murphy, and Adrian E. Raftery. 2016. “Mclust 5: Clustering, Classification and Density Estimation Using Gaussian Finite Mixture Models.” The R Journal 8 (1): 289.

Shah, Kalpit, Charles E. McCormack, and Neil A. Bradbury. 2014. “Do You Know the Sex of Your Cells?” American Journal of Physiology. Cell Physiology 306 (1): C3–18.

Sullivan, K. M., A. Mannucci, C. P. Kimpton, and P. Gill. 1993. “A Rapid and Quantitative DNA Sex Test: Fluorescence-Based PCR Analysis of X-Y Homologous Gene Amelogenin.” BioTechniques 15 (4): 636–38, 640–41.

Tannenbaum, Cara, Danielle Day, and Matera Alliance. 2017. “Age and Sex in Drug Development and Testing for Adults.” Pharmacological Research: The Official Journal of the Italian Pharmacological Society 121 (July): 83–93.

Toker, Lilah, Min Feng, and Paul Pavlidis. 2016. “Whose Sample Is It Anyway? Widespread Misannotation of Samples in Transcriptomics Studies.” F1000Research 5 (August): 2103.

Wang, Zichen, Caroline D. Monteiro, Kathleen M. Jagodnik, Nicolas F. Fernandez, Gregory W. Gundersen, Andrew D. Rouillard, Sherry L. Jenkins, et al. 2016. “Extraction and Analysis of Signatures from the Gene Expression Omnibus by the Crowd.” Nature Communications 7 (September): 12846.

Wishart, David S., Yannick D. Feunang, An C. Guo, Elvis J. Lo, Ana Marcu, Jason R. Grant, Tanvir Sajed, et al. 2018. “DrugBank 5.0: A Major Update to the DrugBank Database for 2018.” Nucleic Acids Research 46 (D1): D1074–82.

Wishart, David S., Craig Knox, An Chi Guo, Savita Shrivastava, Murtaza Hassanali, Paul Stothard, Zhan Chang, and Jennifer Woolsey. 2006. “DrugBank: A Comprehensive Resource for in Silico Drug Discovery and Exploration.” Nucleic Acids Research 34 (Database issue): D668–72.

Woitowich, Nicole C., Annaliese Beery, and Teresa Woodruff. 2020. “Meta-Research: A 10-Year Follow-up Study of Sex Inclusion in the Biological Sciences.” eLife 9: e56344.

Xu, Jin, Xinxin Peng, Yuxin Chen, Yuezheng Zhang, Qin Ma, Liang Liang, Ava C. Carter, Xuemei Lu, and Chung-I Wu. 2017. “Free-Living Human Cells Reconfigure Their Chromosomes in the Evolution back to Uni-Cellularity.” eLife 6 (December). https://doi.org/10.7554/eLife.28070.

Zhou, Baiyu, and Wing Hung Wong. 2011. “A BOOTSTRAP-BASED NON-PARAMETRIC ANOVA METHOD WITH APPLICATIONS TO FACTORIAL MICROARRAY DATA.” Statistica Sinica 21 (2): 495–514.

Zhu, Yuelin, Sean Davis, Robert Stephens, Paul S. Meltzer, and Yidong Chen. 2008. “GEOmetadb: Powerful Alternative Search Engine for the Gene Expression Omnibus.” Bioinformatics 24 (23): 2798–2800.

Zopf, Y., C. Rabe, A. Neubert, K. G. Gassmann, W. Rascher, E. G. Hahn, K. Brune, and H. Dormann. 2008. “Women Encounter ADRs More Often than Do Men.” European Journal of Clinical Pharmacology 64 (10): 999–1004.

