## Supplemental Figures and Table Legends for "Large-Scale Labeling and Assessment of Sex Bias in Publicly Available Expression Data"

### SUPPLEMENTAL TABLES

**Supplementary Table 1.** Sex breakdown by sample (A) and study (B). These data are divided by organism, data type (microarray or RNA-seq), and label type ("metadata" or "expression"), where expression refers to the labels from our model. The sample-level table includes counts of male, female, and unlabeled samples, while the study-level table includes the counts of studies in each of the study types (see Methods 2-1). Fractions of samples and studies are also included.

**Supplementary Table 2.** Accuracy assessment. A) List of assessment datasets for each organism, data_type, and the number of studies and samples and sex breakdown of each dataset. B) Accuracy of sex labeling in each assessment dataset at varying "cutoff" probability scores.

**Supplementary Table 3.** Platform accuracy. A) List of all platforms, the number of samples of that platform, and accuracy and fraction of samples from that platform labeled by our model (at cutoff threshold 0.7). B) Fractions of all samples belonging to platforms with poor accuracy ("poor_accuracy", defined as <70% concordance), not included in the extended test set ("not_in_test"), and included samples after removing poor accuracy or untested platforms ("included"). The approximate accuracy ("approx_accuracy") is calculated by multiplying the accuracy of that platform by the prevalence of the platform overall, and the approximate fraction of samples labeled ("approx_frac_labeled") is calculated by multiplying the fraction labeled in platform a platform by the prevalence of the platform. These refer to the accuracy and fraction labeled in the included set.

**Supplementary Table 4.** Counts by sample source type. The number of samples assigned to each source type.

**Supplementary Table 5.** Y chromosome loss across 87 male cell lines. Includes the cell line name ("cl_name") and accession, number of samples and studies, number of samples labeled male and female, and the median P(male) across all samples ("median_p_male")

**Supplementary Table 6.** Misannotation estimates in (A) single versus mixed sex studies, (B) mixed sex studies by comparison to other expression-based methods, and (C) mixed sex studies based on the distribution of sample sex scores. Estimates are given at varying thresholds of our model (A + B) or for clustering cutoffs (C).

**Supplementary Table 7.** (A) List of drug exposure studies. Each row is a study/drug pair, and includes the DrugBank ID ("dbID"), "drug_name", and ATC code. The "sample_terms" column refers to whether the study contains samples with both control and drug terms ("both") or only drug terms ("drug-only"). (B) Sex bias by drug. Each row contains the organism, drug (with DrugBank ID, name, and ATC code), number of studies, and fractions male, female, and mixed sex studies.

**Supplementary Table 8.** Studies with cancer (A) (ATC class L) and nervous system drugs (B) (ATC class N), organized by drug and study sex breakdown.

### SUPPLEMENTAL FIGURES

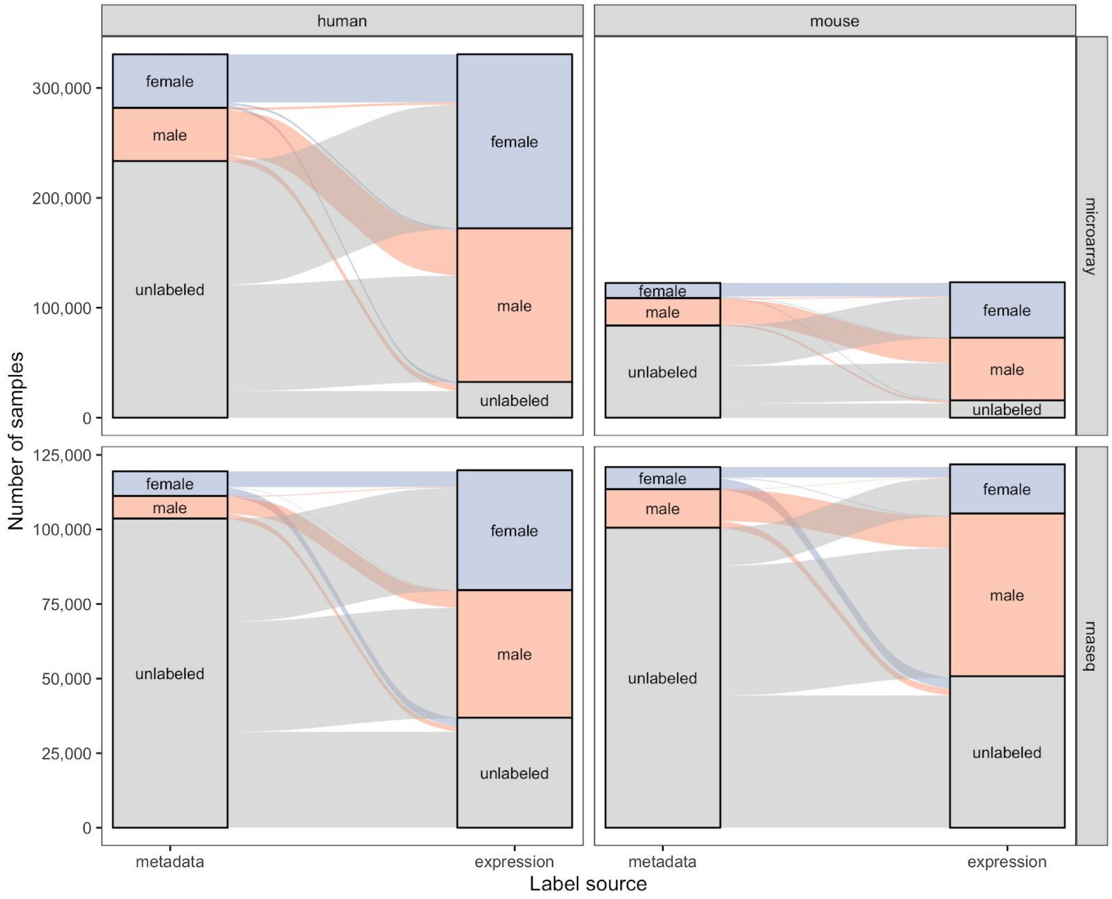


**Supplementary Figure 1.** Sample breakdown. Same as Figure 1 but at a sample rather than study level.

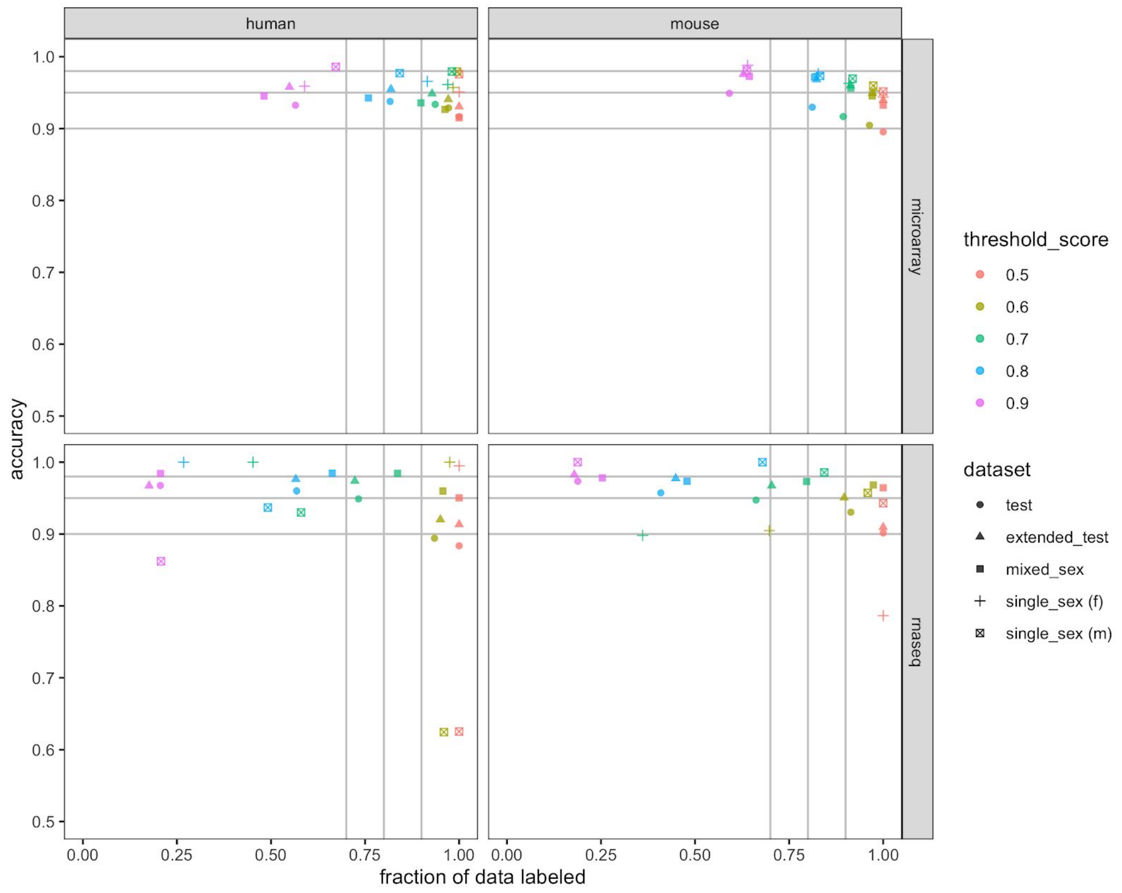


**Supplementary Figure 2.** Accuracy and fraction of data labeled at certain sample sex score cutoffs. Each color is a different sample sex score cutoff, the shape of the point indicates the accuracy and fraction labeled on the dataset it is being evaluated on. Gray lines highlight the 0.9, 0.95, 0.98 accuracy cutoffs and 0.7, 0.8, and 0.9 data fractions.

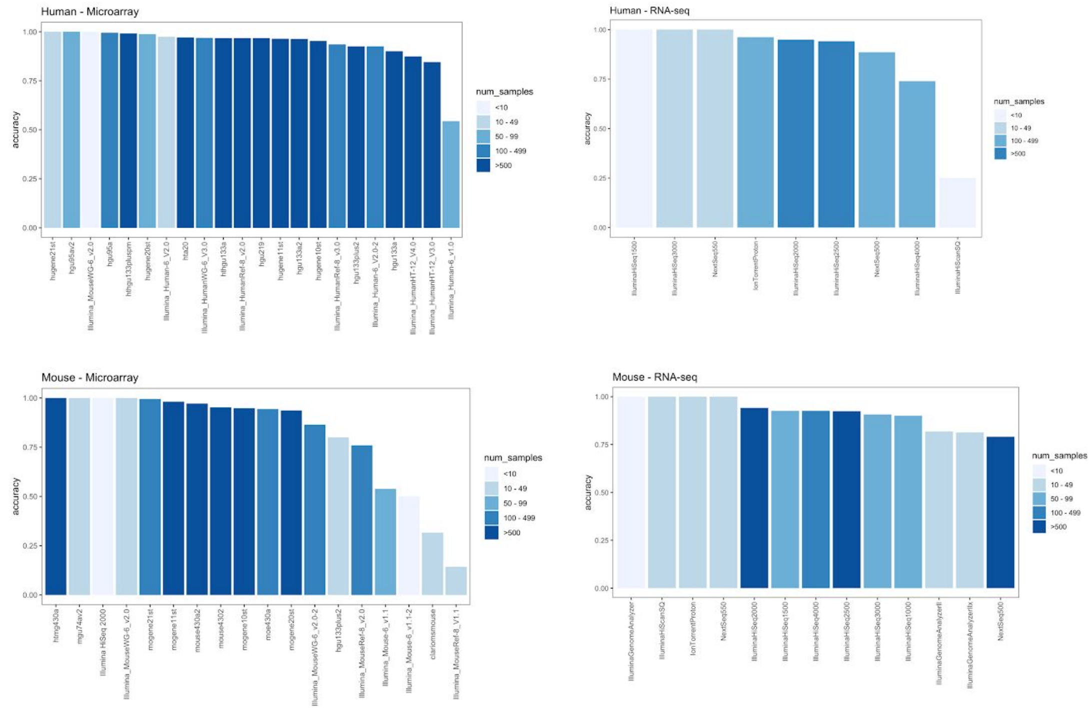


A)

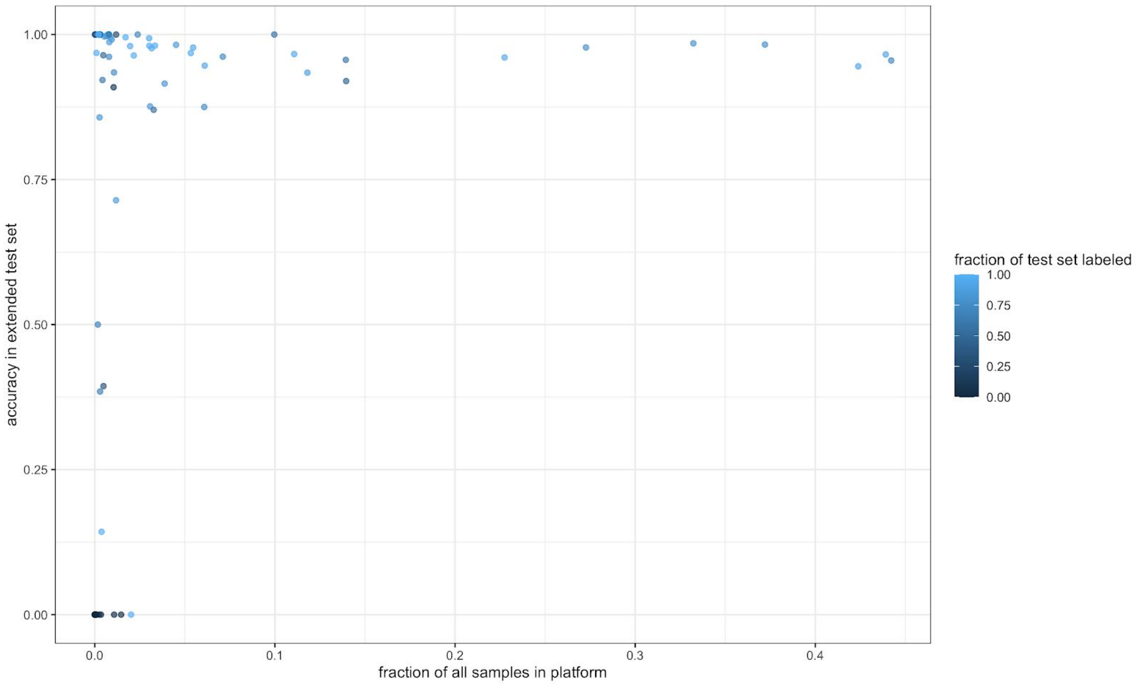


B)

**Supplementary Figure 3.** A) Accuracy across platforms. B) Platform accuracy in the test set versus the fraction of all samples in that platform, where each point is a platform. Bars and points are colored by the number of samples in that platform.

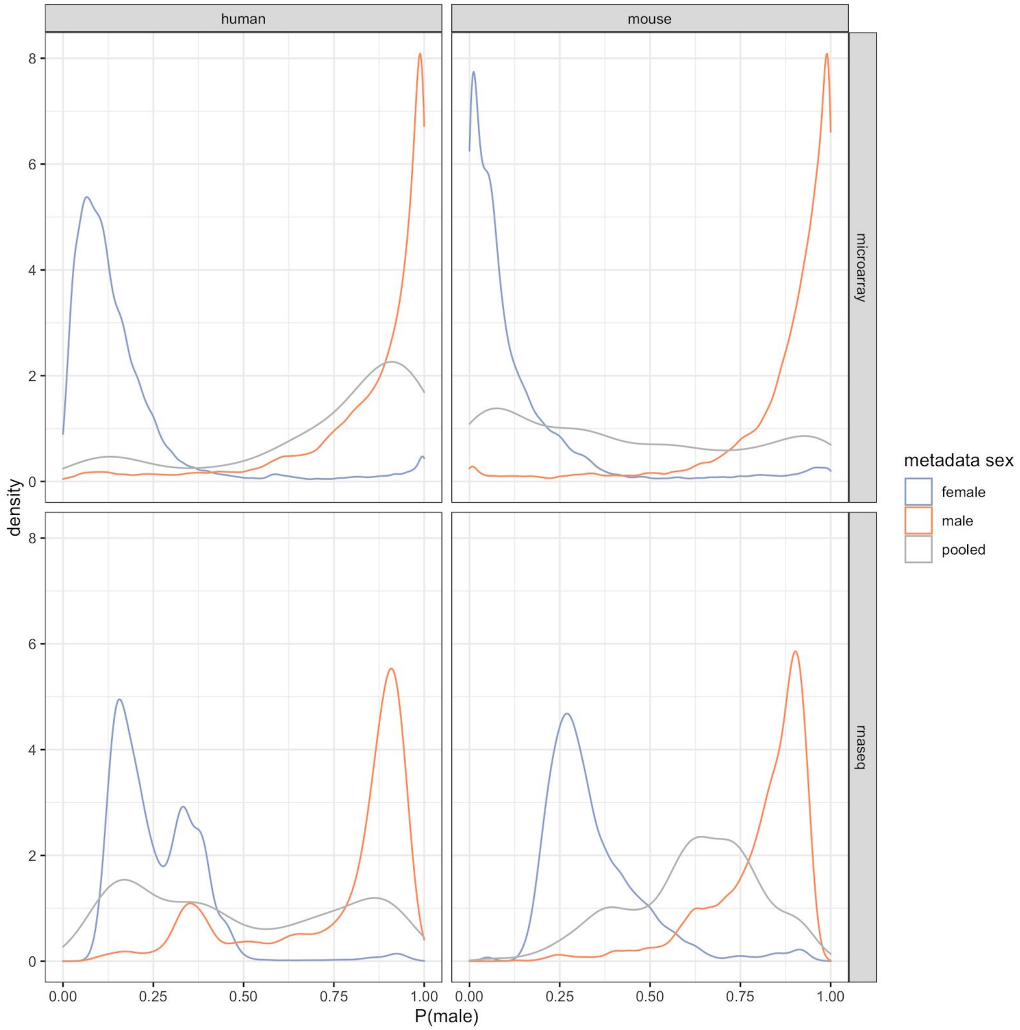


**Supplementary Figure 4.** Sample sex score distributions of female, male, and pooled samples by organism and technology type.

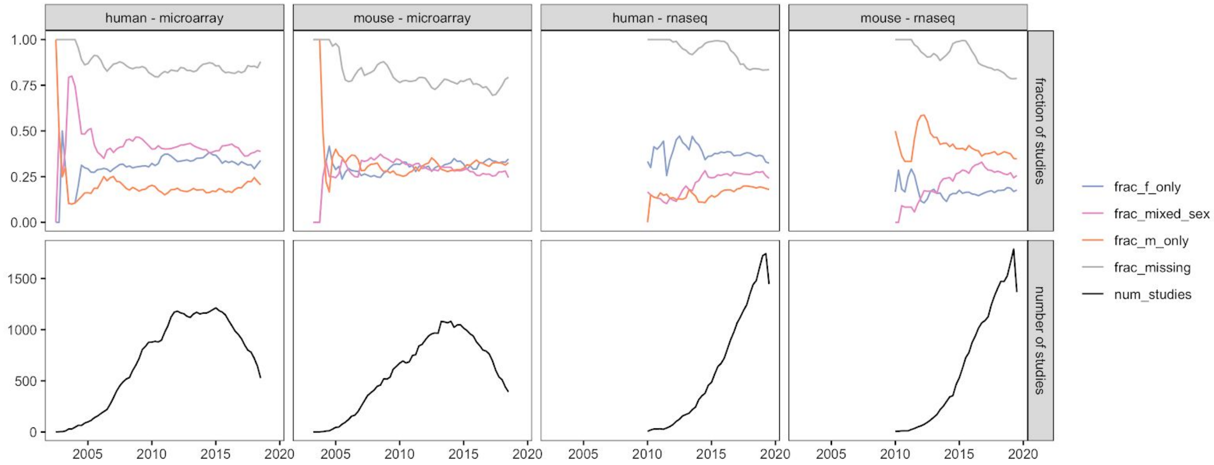

**Supplementary Figure 5.** Sex breakdown over time, by organism and data type. The top panel shows the fraction of studies with either missing sex labels (gray), or imputed female-only (blue), male-only (orange), or mixed sex (pink) samples.  The bottom panel provides the total number of studies over time. Data were smoothed using a three-month window.

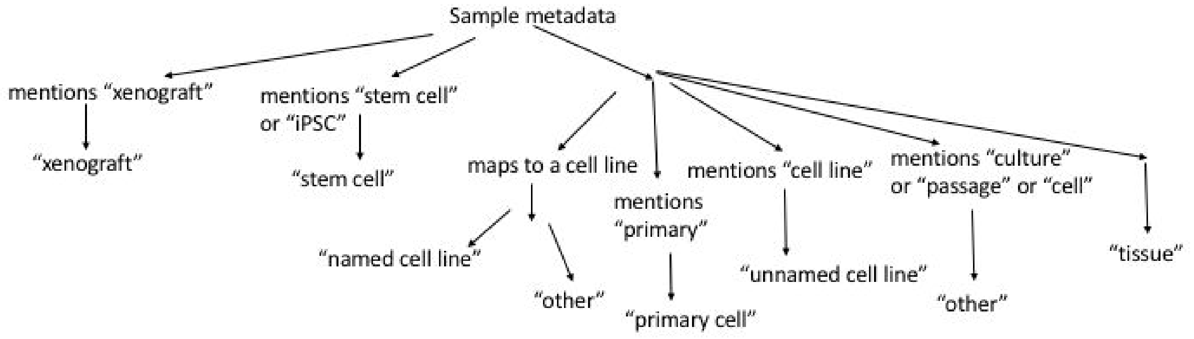


**Supplementary Figure 6.** Sample source assignment. Diagram showing logic of how samples were assigned to a sample source type based on metadata.

A)

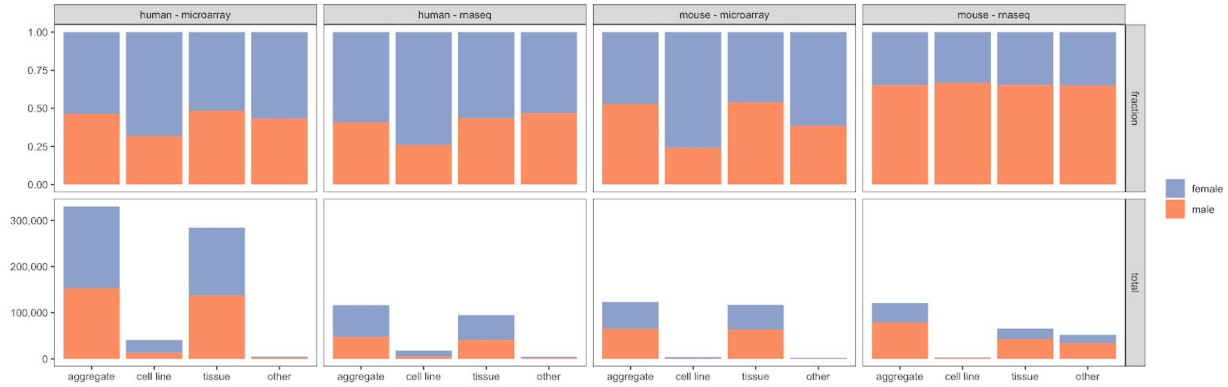


B)

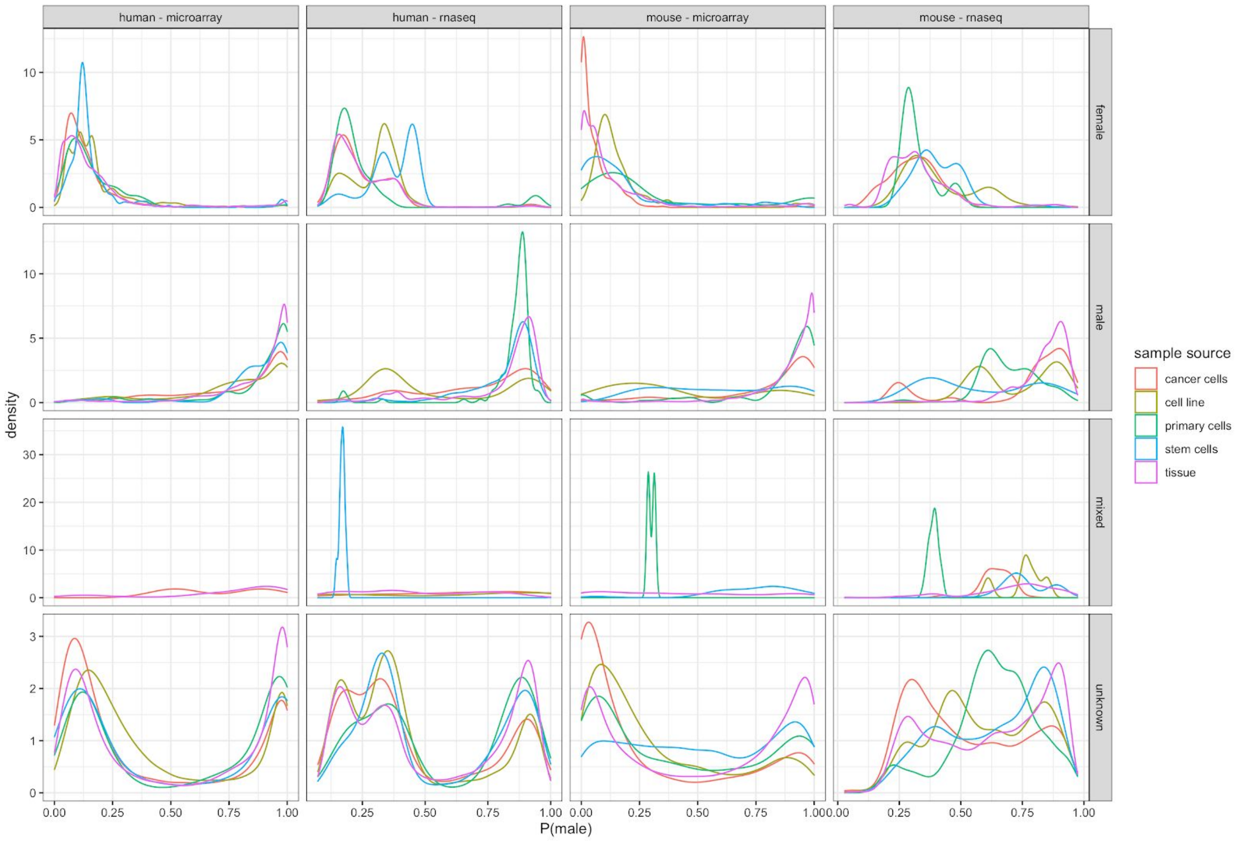


**Supplementary Figure 7.** A) Sex breakdown overall and in cell line versus tissue data. Bars are colored female (dark blue) or male (orange) to indicate the fraction (top row) or count (bottom row) of samples of that sex. B) Distribution of sample sex scores in cancer (red), cell line (yellow), primary cell (green), stem cell (blue), and tissue (purple) samples; separated by metadata sex (rows), and organism/data source (columns).

A)

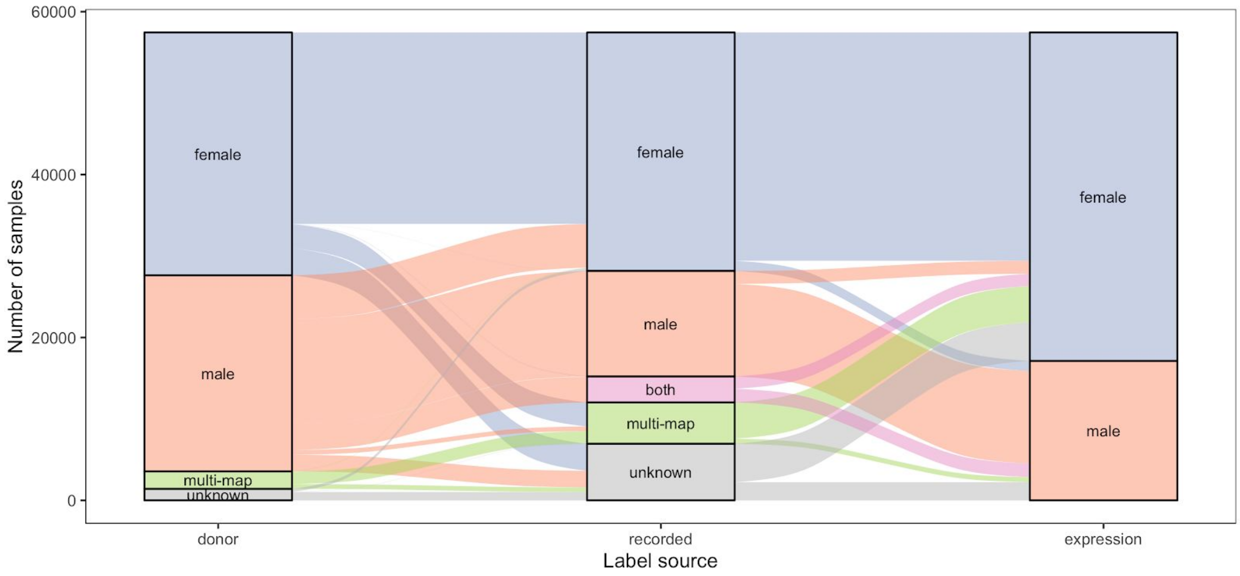


**Supplementary Figure 8.** Alluvial diagram showing cell line sex label “switching” and comparison of expression imputed sex labels to annotations of donor and recorded sex for that cell line.

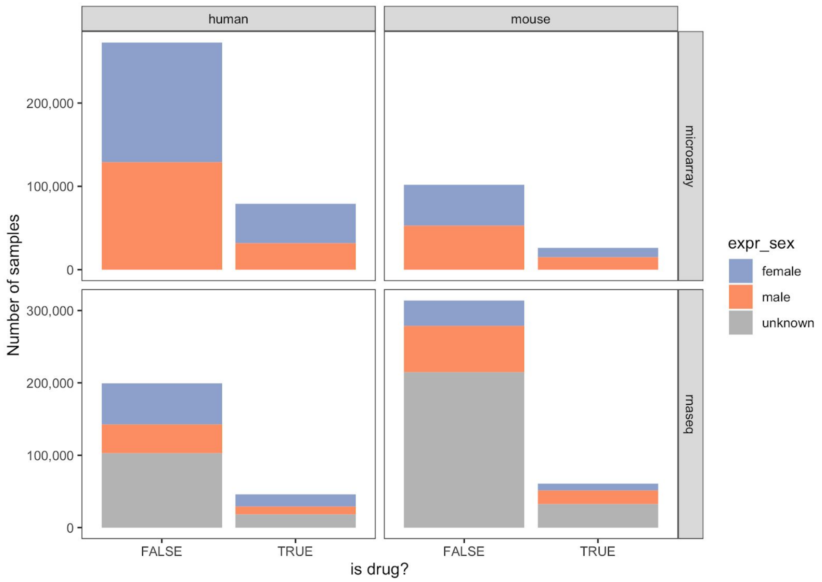


A)

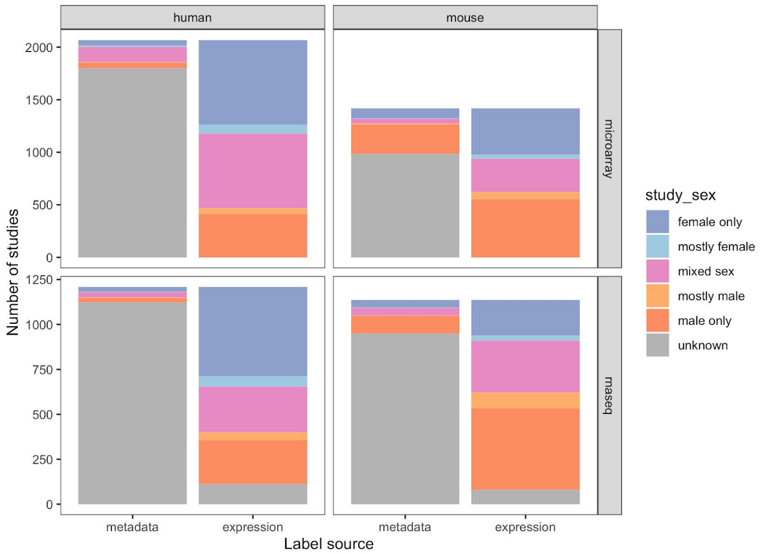


B)

**Supplementary Figure 9.** Sex breakdown of drug samples (A) and studies (B) for *drug mention* data (see Methods 5-1).

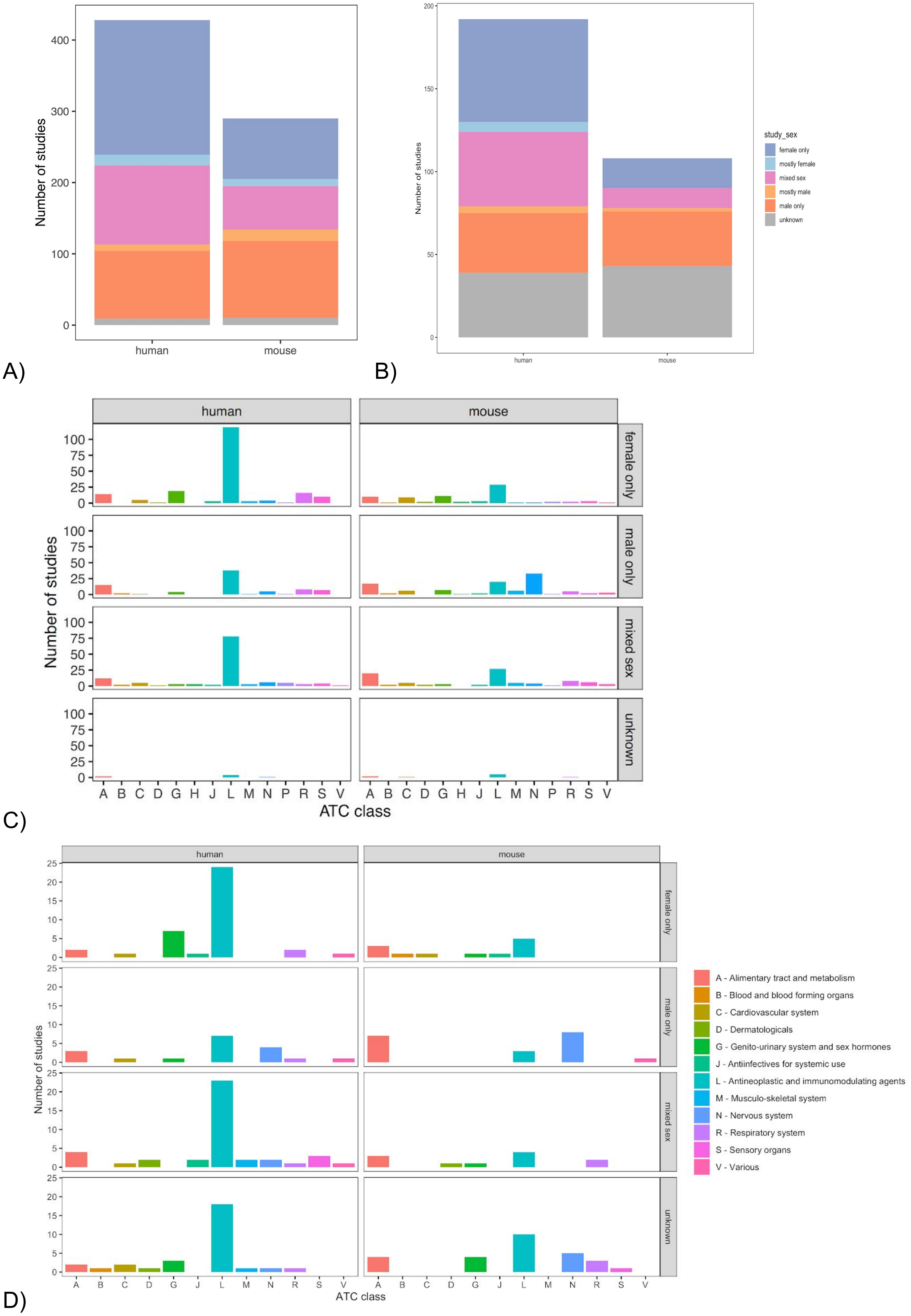

A)
Number of studies
400
300
200
100
0
human
mouse
B)
Number of studies
200
150
100
50
0
human
mouse
study_sex
female only
mostly female
mixed sex
mostly male
male only
unknown
C)
human
mouse
female only
male only
mixed sex
unknown
Number of studies
100
75
50
25
0
A B C D G H J L M N P R S V
ATC class
D)
human
mouse
female only
male only
mixed sex
unknown
Number of studies
25
20
15
10
5
0
A B C D G J L M N R S V
ATC class
A - Alimentary tract and metabolism
B - Blood and blood forming organs
C - Cardiovascular system
D - Dermatologicals
G - Genito-urinary system and sex hormones
J - Antiinfectives for systemic use
L - Antineoplastic and immunomodulating agents
M - Musculo-skeletal system
N - Nervous system
R - Respiratory system
S - Sensory organs
V - Various

**Supplementary Figure 10.** Sex breakdown of samples directly mapping to drugs (*drug exposure studies* - see Methods 5-1) and manually curated drug data from Wang et al. (Wang et al. 2016) matches patterns seen in drug breakdown data in terms of overall study sex breakdown (A,B) and ATC class breakdown (C, D for drug exposure studies and Wang et al respectively).
